# 1-Aminocyclopropane-1-carboxylic acid oxidase determines the fate of ethylene biosynthesis in a tissue-specific way to fine-tune development and stress resilience

**DOI:** 10.1101/2024.02.01.578397

**Authors:** M. Houben, J. Vaughan-Hirsch, J. Pattyn, W. Mou, S. Roden, A. Martinez Roig, E. Kabak, S. Rodrigues, A. Polko, B. De Coninck, J. J. Kieber, A. Voet, B. Van de Poel

**Affiliations:** Division of Crop Biotechnics, Department of Biosystems, University of Leuven, Willem de Croylaan 42, 3001 Leuven, Belgium; College of Life and Environmental Sciences, Hangzhou Normal University, Hangzhou, China; Division of Biochemistry, Molecular and Structural Biology, Department of Chemistry, University of Leuven, 3001 Leuven, Belgium; Department of Biology, University of North Carolina, Chapel Hill, NC, 27599, USA; KU Leuven Plant Institute (LPI), University of Leuven, Kasteelpark Arenberg 31, 3001 Leuven, Belgium

**Keywords:** ACC-oxidase, ethylene, vegetative development, generative development, abiotic stress, biotic stress

## Abstract

Ethylene is involved in several developmental processes and responses towards (a)biotic stress. In seed plants, ethylene is produced from its precursor 1-aminocyclopropane-1-carboxylic acid (ACC) by the enzyme ACC-oxidase (ACO). Despite its key role in ethylene synthesis, the *ACO* gene family has not yet been fully characterized. We investigated the five *ACO* members of *Arabidopsis thaliana* and revealed a tissue-and developmentally specific expression pattern. Furthermore, the five ACO enzymes each have a different capacity to produce ethylene. Combined, this allows for a precise spatial and temporal regulation of ethylene synthesis. At the sub-cellular level, we uncovered that ACOs reside in the cytosol, where ethylene is likely synthesized, but surprisingly also in the nucleus. Using reverse genetics of single and higher order *aco* mutants we revealed a high degree of gene redundancy and minimal phenotypes. A lack of ethylene synthesis by knocking out all five *ACOs* did not impair normal vegetative and generative development but did influence processes associated with high rates of ethylene production such as petal abscission. This suggests that ethylene is not a prime regulator of development, but more a moderator. We also showed that the inability to synthesize ethylene impairs some abiotic (nutrient deficiency and metal toxicity) and biotic (*Botrytis cinerea*) stress responses, similar as plants insensitive towards ethylene, corroborating the role of ethylene in mediating stress responses. In conclusion, the *ACO* gene family enables plants to fine-tune their ethylene synthesis rates, but a lack their off is not crucial for normal development and stress survival.

## Introduction

The volatile hydrocarbon ethylene is produced by plants and other living organisms such as certain fungi and bacteria (Van de Poel *et al*., 2014*a*; Papon and Binder, 2019). In plants, ethylene is an important gaseous hormone, involved in growth, development, and stress responses. For example, in seed plants, ethylene is involved in germination, vegetative development, climacteric fruit ripening, abscission, senescence and much more (Abeles *et al*., 1992). Furthermore, ethylene is also linked with both biotic and abiotic stress responses (Chen *et al*., 2022). Many ethylene-regulated processes are of great agricultural importance (Chang, 2016).

In seed plants, ethylene is produced by a dedicated two-step pathway. First, S-adenosyl-L-methionine is cleaved into 1-aminocyclopropane-1-carboxylic acid (ACC) and 5’-methylthioadenosine by the enzyme ACC synthase (ACS). Subsequently, ACC is converted into ethylene gas by ACC oxidase (ACO) (Pattyn et al., 2021; and references therein). ACO is a dedicated enzyme conserved in seed plants, while homologs are not retrieved in genomes of non-seed plants, suggesting the ACC-route of ethylene biosynthesis is present only in seed plants (Li *et al*., 2018, 2022*b*; Jiao *et al*., 2020; Van de Poel & de Vries., 2023). Some ethylene-producing fungi seem to have a functional ACO-like homolog, which could be related to the plant ACOs (Li *et al*., 2023). Contrary, *ACS* genes are retrieved in all plants and some algal genomes (Li *et al*., 2018; Jiao *et al*., 2020), indicating ACC synthesis is more ancient than its effective conversion into ethylene by ACO (Li *et al*., 2022*c*). In Arabidopsis, ACC is synthesized by a group of eight functional PLP-dependent ACS proteins, that can form homo-and heterodimeric complexes (Boller *et al*., 1979; Tsuchisaka *et al*., 2009). Furthermore, ACS activity is determined by its post-translational stability controlled by the differential phosphorylation of the type-specific C-terminal regulatory domain (Park *et al*., 2021). Besides the regulation of ACC synthesis, ACC homeostasis is also controlled by ACC conjugation and transport (Van de Poel and Van Der Straeten, 2014). ACC can be sequestered through conjugation into malonyl-ACC (MACC; Amrhein *et al*., 1981; Hoffman *et al*., 1982), ɣ-glutamyl-ACC (GACC; Martin *et al*., 1995) and jasmonyl-ACC (JA-ACC; Staswick and Tiryaki, 2004). Furthermore, ACC is mobilized in plants through amino acid transporters Lysine-Histidine Transporter 1 (LHT1; Shin *et al*., 2015; Li *et al*., 2022*a*) and LHT2 (Choi *et al*., 2019).

Based on the observation that feeding ACC to plant tissue raises ethylene production, it was postulated that ACS is the rate limiting step in ethylene synthesis (Yang and Hoffman, 1984), but newer studies found that ACO can also be rate limiting during certain developmental stages or stress conditions (Houben and Van de Poel, 2019). In Arabidopsis, ACO is encoded by a group of five genes (Houben and Van de Poel, 2019), that belong to the 2-oxoglutarate-dependent dioxygenases (2OGD) superfamily requiring iron for catalytic activity (Bouzayen *et al*., 1991). Furthermore, ACO requires ascorbate (vitamin C) as reductant (Ververidis and John, 1991; Murphy *et al*., 2014) and bicarbonate and molecular oxygen as activators for catalytic activity (Peiser *et al*., 1984; Dilley *et al*., 2013). Besides ethylene, ACO also produces CO_2_ and a cyanide ion (CN^-^), which is rapidly detoxified into β-cyanoalanine (Peiser *et al*., 1984). ACO is presumed to be localized in the cytosol (Peck *et al*., 1992; Reinhardt *et al*., 1994; Chung *et al*., 2002; Hudgins *et al*., 2006; Kawai *et al*., 2014; Tu *et al*., 2019), contesting other studies that reported ACO to be membrane-associated (Kende, 1989; Rombaldi *et al*., 1994; Ramassamy *et al*., 1998). ACO is primarily regulated transcriptionally with only few transcription factors identified (Houben and Van de Poel, 2019). Some studies reported redox-mediated control of ACO protein stability and activity through cysteine modifications (Dixon *et al*., 2005; Aroca *et al*., 2015; Datta *et al*., 2015; Jia *et al*., 2018; Fournier *et al*., 2019; Tachon *et al*., 2019; Liu *et al*., 2023), revealing a novel layer of ethylene control.

Despite the discovery of ACO more than 3 decades ago in tomato (Hamilton *et al*., 1990), little is known about the characterization of the *ACO* gene family at the genetic, molecular and biochemical level. Many questions about the ACO isoform-specific biological significance remain. Here we provide a comprehensive characterization of the *ACO* gene family in *Arabidopsis thaliana*. We reveal that ACOs reside subcellular both in the cytosol and nucleus using transcriptional and translational reporter lines. We also demonstrate that the different ACOs have distinct enzyme kinetics. A functional characterization of single and higher order *aco* mutants showed a high degree of gene redundancy and exposes that ethylene biosynthesis by ACO is not crucial for normal vegetative and generative development and stress responses, but rather is a moderator of these processes.

## Materials and Methods

### Plant material

*Arabidopsis thaliana* ecotype Col-0 was used as wild-type control plants. The individual *aco* T-DNA insertion mutants were obtained from the Nottingham Arabidopsis Stock Centre (NASC) (Supplemental Table S1). Higher order *aco* mutants were obtained by crossing. The ET (ethylene) free lines 1 and 2 (ET free 1 & 2) were obtained from Dr. Li-Jia Qu (Peking University, China). See the Supplementary Methods for growth conditions.

### Generation of CRISPR/Cas9 mutants and ACO reporter lines

The *aco* quintuple lines made in this study (ET free 3 and 4) were created by knocking out *ACO5* using CRISPR/Cas9 in the *aco1aco2aco3aco4* T-DNA quadruple line (obtained by crossing). Two gRNA’s (Supplemental Table S2) were cloned into the *p00178* vector using Goldengate, according to the TSKO protocol (Decaestecker *et al*., 2019). For the creation of the transcriptional *ACO* reporter lines, the promoter region (1807 - 2769 bases upstream from the ATG start codon) of each *ACO* gene was amplified See the Supplementary Methods for the cloning and transformation steps.

### Genotyping and quantitative real-time PCR

Genotyping was done using T-DNA and gene-specific primers (Supplemental Table S2). CRISPR mutants were genotyped by PCR and sequencing. The expression of individual *ACO* genes in the single *aco* T-DNA mutants was validated using RT-qPCR (4 replicates). See qPCR details in the Supplementary Methods.

### Phenotyping

For dark-grown phenotyping, seedlings were grown on horizontal plates for 4 days in the dark at 21 °C. Light-grown seedlings were cultivated on vertical plates for 10 days. Hypocotyl and root length of individual seedlings were measured using ImageJ. Rosette diameter was measured of 3-week-old soil-grown plants using ImageJ. The number, length and branching of inflorescences were recorded from 4-week-old soil-grown plants. Floral organs, silique length and seed size were observed using a stereomicroscope (Olympus SZX) and quantified using ImageJ. Leaf senescence rate was assessed in 52-day-old soil-grown plants by quantifying leaves showing signs of senescence.

### GUS staining and fluorescence microscopy

For imaging reporter lines after ethylene/ACC treatment, 5-day-old plate-grown seedlings were incubated with 1 ppm ethylene, or 1 - 5 µM ACC (as a proxy for ethylene), for 4 hours before GUS staining (see Supplementary Methods).

To visualize GFP in transcriptional reporters, live plants were mounted in water and imaged on a Leica Thunder Imager, using the brightfield and GFP filter cube (excitation 450-490 nm, emission 500-550 nm). To visualize the ACO translational reporter, live plants were stained with propidium iodide (PI; 10 µg/mL in water) and DAPI (1 µg/mL in water) for 5 minutes each, washed briefly in water and imaged immediately on a Leica SP8 confocal microscope. For imaging cleared plants, tissue was first fixed in 4 % formaldehyde (v/v) in PBS for 1 hour, then washed in PBS and incubated with ClearSee for at least 1 day (Ursache *et al*., 2018). Tissues was then stained with DAPI (1 µg/mL in ClearSee) for 30 minutes and washed in ClearSee for 1 hour before imaging on a Leica SP8 confocal microscope. Sequential scanning was used with an excitation laser at 488 nm (for GFP), 535 nm (for PI) and 405 nm (for DAPI), and fluorescence detectors at 493-569 nm (for GFP), 541-651 nm (for PI) and 497-565 nm (for DAPI).

### Ethylene, ACC and MACC quantification

Ethylene production was measured of dark-grown 4-day-old seedlings in airtight GC-vials (10 mL) with solid MS medium (0.8 % agar w/v) with or without ACC. For light-grown seedlings, they were first grown on MS media supplemented with ACC (0.5 µM and 5 µM) and subsequently transferred to 4 mL vials with 1 mL liquid MS supplemented with the same concentrations of ACC and incubated for 48 h. Next, 1 mL of headspace was sampled with a syringe and injected into a Shimadzu gas chromatograph (GC-2010). The GC was equipped with a capillary carboxen 1010 PLOT column (30 m x 0.53 mm) with N_2_ (35 mL/min) as carrier gas. The injector and column temperature were set at 200 °C and 40 °C respectively. Ethylene was injected automatically by the PAL AOC6000 pipetting robot using a split-method (2/1) and detected using a flame ionization detector at a temperature of 250 °C. Following ethylene measurements, seedlings were weighed and ethylene production by mass was calculated.

ACC and MACC were detected by the Lizada & Yang method (Lizada and Yang, 1979) as described by Bulens et al. (2011) and explained in the Supplementary Methods.

### Recombinant ACO *in vitro* activity assay

The coding sequence of each *ACO* was amplified and cloned into the *pET28a* bacterial expression vector and transformed into E. coli strain BL21 (DE3). Starter cultures were grown in 5 mL LB media with 50 μg.mL^-1^ kanamycin and chloramphenicol overnight at 37 °C shaking at 200 rpm. Next, 2 L cultures were inoculated and grown at 37 °C shaking at 100 rpm until an OD600 of 0.8 was reached. Cultures were cooled on ice for 30 min, protein expression induced by adding isopropyl ß-D-1-thiogalactopyranoside (IPTG) to a final concentration of 0.5 mM, and then grown overnight at 18 °C shaking at 100 rpm. See the Supplementary Method for ACO protein purification.

Recombinant *in vitro* ACO activity measurements were done as described by (Van de Poel *et al*., 2014*b*). Briefly, 5 μg of purified ACO protein was incubated in freshly prepared activity buffer containing 50 mM MOPS, 5 mM ascorbic acid, 20 mM sodium bicarbonate, 10 % glycerol (v/v), 0.1 mM DTT and ACC as substrate. The reaction was incubated in 4 mL airtight GC vials for 60 min at 30 °C while shaking. One mL of headspace was sampled and analyzed for ethylene content using GC as described above.

### Abiotic stress experiments

For abiotic stress experiments of plate-grown plants, seedlings were sterilized and grown on plates containing the indicated concentrations of NaCl for salt stress and diluted MS medium for nutrient deficiency stress. After 10 days, plates were imaged, and root length was measured using ImageJ. For heavy metal stress, 5-day-old seedlings were transferred to plates containing indicated concentrations of heavy metals and grown for another 10 days. Images were taken of the plates at 6 and 10 days to allow measurements of primary root length (Remy and Duque, 2016).

For drought and salt stress experiments using soil-grown plants, 4 plants per pot were grown with equal watering before the start of the treatment. For drought experiments, water was withheld after 3 weeks for 3 weeks. Plants were then rewatered and after 1 week recovery and survival rates were determined (Yang *et al*., 2021). For salt stress experiments, 3-week-old plants were watered with 1 L NaCl solution per tray at the indicated concentrations. Plants were then grown for 10 days before scoring survival rates, with non-saline water being given during the recovery period (Jiang *et al*., 2013). For submergence experiments of soil-grown plants, one plant per pot was grown for 3 weeks before submergence. Plants were submerged in the dark for 3 days. The water was previously equilibrated to room temperature, and plants were submerged 5 cm above the rosettes. After 3 days, plants were removed and returned to the growth room, and survival rates were scored after 7 days recovery (Schmidt *et al*., 2018). Images were taken before and immediately after each abiotic stress period, and after the recovery period.

### Biotic stress experiments with *Botrytis cinerea*

Four-week-old plants were tested for their susceptibility towards infection by *B. cinerea* prior to flowering. Briefly, *B. cinerea* spores (see the Supplementary Methods of sporulation induction) were taken out of the −80 °C and diluted to a concentration of 5 × 10^5^ spores/mL in 0.5 x potato dextrose broth. Five µL of diluted spores were applied to three leaves per plant for at least 20 plants. The plants were arranged in trays following a randomized block design and a cover was placed over the trays to simulate high humidity conditions. Lesion sizes were measured (n > 50) at three-and four-days post inoculation.

### Statistical analysis

Statistical analysis was performed, and graphs were made using Graphpad Prism 8.4. For all *aco* mutant phenotyping, outlier analysis was performed using the ROUT method with default settings (Q = 1 %), and outliers removed before further analysis. Normal distribution (D’Agostino and Pearson test) and equal variance (Brown-Forsythe test) was tested. For samples normally distributed with non-equal variance, Welch ANOVA and Games-Howell’s post hoc test was performed. For samples non-normally distributed, the non-parametric Kruskall-Wallis and Dunn’s post-hoc test was performed. The tests used are indicated in the figure legends.

## Results

### ACO dictates ethylene production levels of Arabidopsis seedlings in a redundant manner

To study the role of the *ACO* gene family in regulating ethylene biosynthesis, plant development and (a)biotic stress resilience, we first developed double *aco* mutants by crossing the available single T-DNA insert *aco* mutants (Supplemental Figure S1A). The expression level of the individual *ACO* genes in the single mutant background was validated by means of RT-qPCR in light-grown seedlings and is presented in Supplemental Figure S1B. Subsequently, a higher order triple *aco1,3,4* and quadruple *aco1,2,3,4* was made by crossing. Finally, a quintuple *aco1,2,3,4,5* mutant was generated by knocking out *ACO5* using CRISPR/Cas9 technology (Supplemental Figure S2) in the quadruple *aco1,2,3,4* T-DNA mutant. For this quintuple mutant, 2 lines (ET free 3 and ET free 4) were generated and compared to the recently published ET free lines (ET free 1 and ET free 2) which are CRISPR/Cas9 knock-out mutants of all five *ACOs* and unable to produce detectable levels of ethylene (Li *et al*., 2021).

To infer the role of individual ACOs on Arabidopsis ethylene production, we measured ethylene production of 4-day-old dark-grown seedlings with and without feeding 2 µM ACC (Figure 1A). In the absence of ACC feeding, ethylene emission rates were close to or below the detection limit for both the wild type and all *aco* mutants. In the presence of ACC, the single *aco* mutants, showed similar ethylene production levels as the wild type, indicating that there is gene redundancy amongst the *ACO* family. For the double *aco* mutants, ethylene levels were slightly reduced but not always significant. The quadruple *aco* mutant produced less ethylene, while the *aco* quintuple mutants (both the previously published ET free lines ET free 1 and ET free 2 (Li *et al*., 2021), and our newly generated *aco* quintuple mutants ET free 3 and ET free 4) did not produce any detectable levels of ethylene, confirming their inability to convert ACC into ethylene. Ethylene production was also measured in 9-day-old light-grown seedlings after 0.5 µM or 5 µM ACC treatment, which confirmed the inability of the *aco* quintuple lines to produce ethylene via ACC (Supplemental Figure S3).

**Figure 1:**
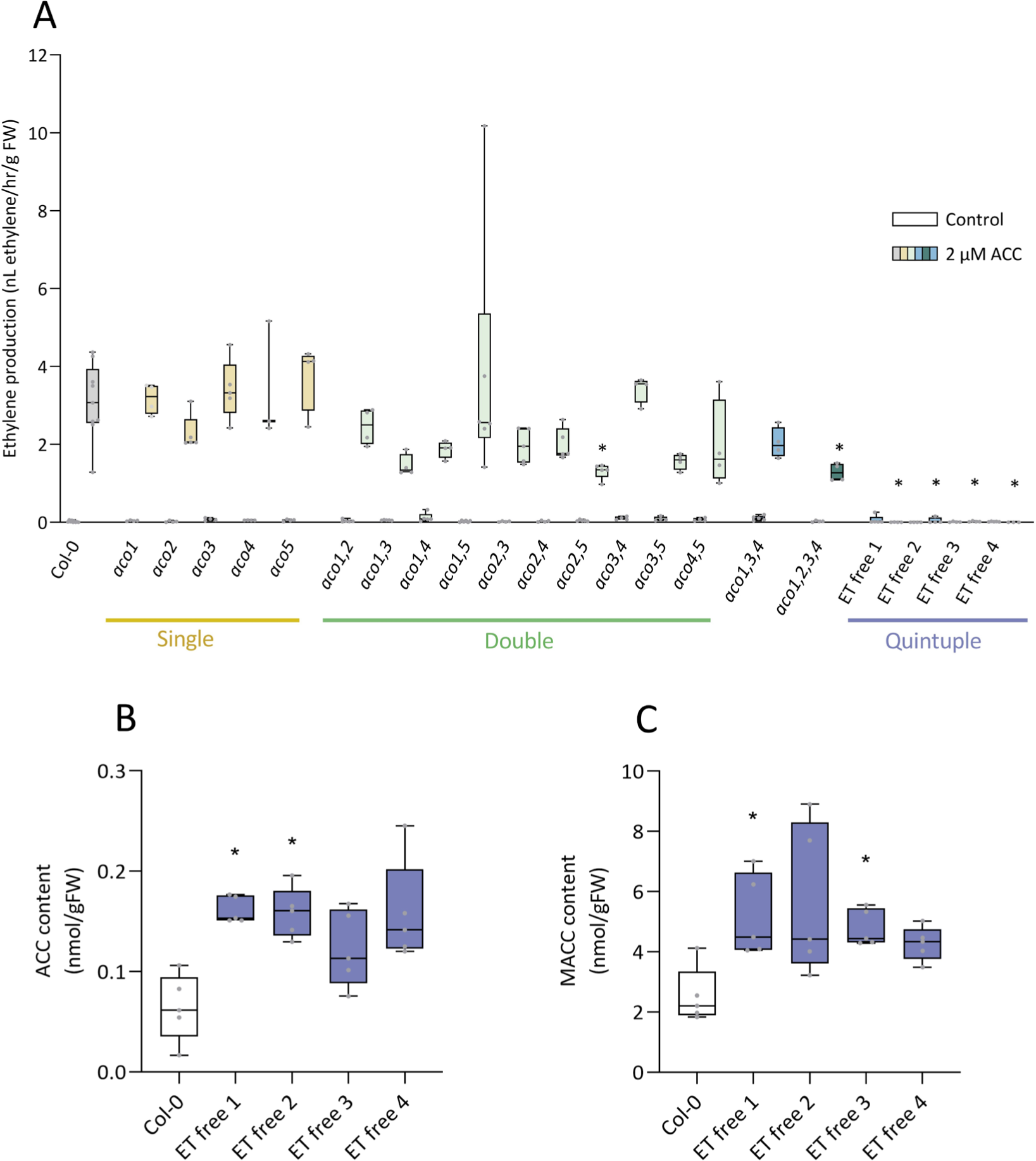
Lack of ACO prevents ethylene production and leads to an accumulation of ACC and MACC. (A) Ethylene production rates of four-day-old dark-grown wild type and *aco* mutant seedlings (n=5) in the presence and absence of 2 µM ACC. (B) Levels of ACC and (C) its conjugate MACC in four-day-old dark-grown seedlings (n=5) of the wild type and *aco* quintuple (ET free 1-4) mutants. Kruskal-Wallis and Dunn’s post-hoc test were performed. Asterisks indicate the significant differences between the wild type and the *aco* mutants (p<0.05).

To unravel if a lack of ethylene production also influences ACC homeostasis, we measured both ACC and MACC levels in the wild type and the quintuple *aco* mutants. Interestingly, all ET-free lines contained higher levels of ACC and its primary conjugate MACC (Figure 1B-C), suggesting a lack of ACC consumption by ACO leads to an accumulation of ACC and its conjugate MACC. This also makes us wonder if certain ACC-specific phenotypes will arise in the quintuple *aco* plants unable to produce ethylene.

### Different ACO isoforms have distinct ACC kinetics

The genetic redundancy for ethylene synthesis made us wonder if individual ACO enzymes have a different capacity to produce ethylene. Therefore, we ectopically expressed the five *ACOs* in E. coli and purified the individual enzymes (Supplemental Figure S4). Next, we assessed the ACC substrate affinity using a dose-response *in vitro* activity assay. We observed that all ACO enzymes were able to produce ethylene from ACC, albeit with a different affinity (Figure 2). ACO1 exhibited the highest enzyme activity (V_max,ACO1_ = 4.67 nmol ethylene/nmol ACO ± 0.26) but a rather low ACC affinity (K_m,ACO1_ = 33.33 μM ± 14.82). Similar to ACO1, ACO3 and ACO4 also had a low affinity (K_m,ACO3_ = 72.67 μM ± 14.67; K_m,ACO4_ = 59.2 μM ± 9.33), but with a maximal activity that is 3-5 times lower (V_max,ACO3_ = 1.91 nmol ethylene/nmol ACO ± 0.12; V_max,ACO4_ = 1.14 nmol ethylene/nmol ACO ± 0.05). Interestingly, ACO2 had the highest affinity (K_m,ACO2_ = 14.82 μM ± 8.25), but the lowest maximal activity (V_max,ACO2_ = 0.15 nmol ethylene/nmol ACO ± 0.02), while ACO5 had the lowest affinity (K_m,ACO5_ = 197.78 μM ± 57.11), but also a low maximal activity (V_max,ACO5_ = 0.58 nmol ethylene/nmol ACO ± 0.07). This data suggests that ACO2 is more effective in producing ethylene in tissues with low ACC levels, while ACO1 is probably used to produce very high levels of ethylene when ample ACC is available.

**Figure 2:**
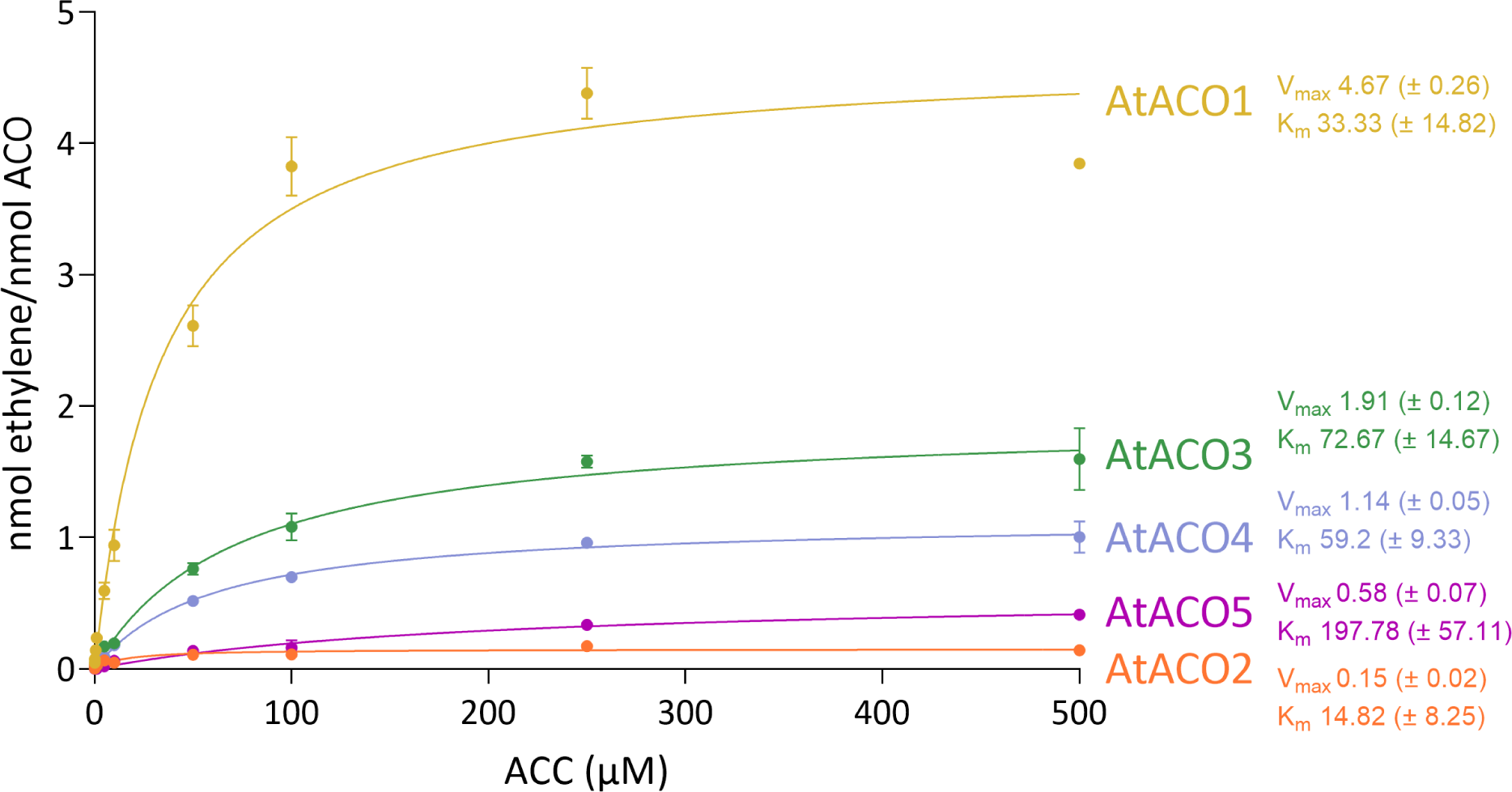
*In vitro* ACO enzyme kinetics show a difference in ACC affinity. Purified recombinant ACO1-5 enzymes were tested for *in vitro* ethylene production capacity when fed with different concentrations of substrate (ACC levels between 0-500 µM). Substrate affinity curves were fitted with a nonlinear regression – Michaelis Menten model and the Vmax and Km values were calculated using Graphpad prism.

### *ACOs* display distinct tissue-specific expression patterns under light and dark conditions

If different ACOs have different affinities for ACC and have different *in vitro* enzyme activities to make ethylene, it made us wonder if there are also differences *in planta*. Therefore, we analyzed the transcriptional (GUS-GFP) and translational (GFP) reporter lines of all five ACOs. We first looked at tissue specific *ACO* expression patterns of light-and dark-grown seedlings. *ACO1* expression is mainly localized the root-shoot junction in light-grown seedlings, and in the root tips of both light-and dark-grown seedlings (Figure 3). Interestingly, *ACO1* expression also seems highly localized to the regions surrounding the root and shoot apical meristems (Supplemental Figure S5). *ACO2* expression was weaker and seems confined to the phloem of leaves and roots of light-grown seedlings (Supplemental Figure S6). *ACO3* is also primarily confined to vascular tissues but is weakly expressed in the hypocotyl of light-grown seedlings and the root vascular tissue of dark-grown seedlings. *ACO4* expression is restricted to the root tip, the root-shoot junction and around meristem regions similar as ACO1, while *ACO5* has a broader expression domain in the root of both light-and dark-grown seedlings and the hypocotyl of dark-grown seedlings. The expression patterns we observed are largely supported by publicly available scRNA-seq data (Supplemental Figure S7).

**Figure 3:**
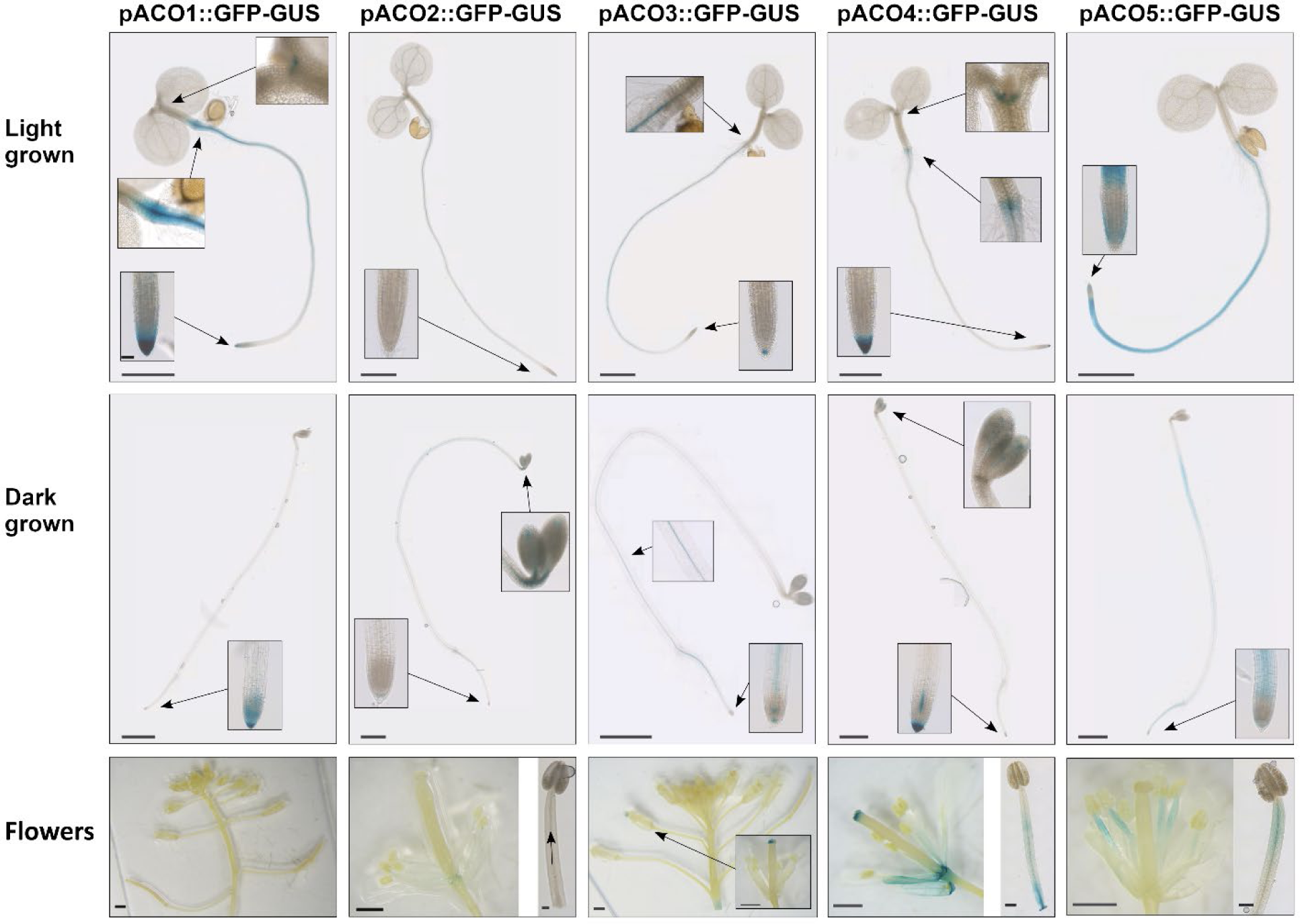
The *ACOs* show distinct tissue-specific expression patterns in seedlings and flowers. Transcriptional p*ACO1-5*:*GUS-GFP* reporters lines were GUS stained following light or dark growth for 5 days and for flower organs of inflorescence. Scale bars are 1 mm for whole seedling images and 50 µm for seedling inlet images. Scale bars are 5 mm for inflorescences and 100 µm for stamen.

We also investigated *ACO* expression patterns in adult and flowering plants. In rosettes, we only observed a GUS signal for the *ACO1* and *ACO4* reporter mainly confined to the leaf petioles and partially to the leaf blade (Supplemental Figure S8). We also observe a differential *ACO* expression in different flowering tissues (Figure 3). *ACO1* was not expressed in any floral tissues. *ACO2* expression is limited to the vascular tissues of the stamen filament, whereas *ACO3* is solely expressed at the stigma of the apical flower of the inflorescence. *ACO4* is more strongly expressed in the stigma, and at the base of all floral organs, including the developing silique. *ACO5* expression is localized specifically to the upper regions of the stamen filaments, with strongest expression in the vascular tissues.

### Translational reporter lines reveal ACO localization to the cytosol and nucleus

Besides the tissue-specific expression, we also wondered what would be the subcellular localization of ethylene biosynthesis, especially because literature has contesting data attributing ACO localization to either the cytosol (Peck *et al*., 1992; Reinhardt *et al*., 1994; Chung *et al*., 2002; Hudgins *et al*., 2006; Kawai *et al*., 2014; Tu *et al*., 2019) or the plasma membrane (Kende, 1989; Rombaldi *et al*., 1994; Ramassamy, 1998). Despite these previous attempts of ACO localization, none of them used stable transformants. Our ACO translational GFP reporter lines allowed us to pinpoint ACO localization to both the cytosol and nucleus of root columella and lateral root cap cells (Figure 4). ACO2 and ACO3 were specifically localized to the phloem of the root vascular bundle (Figure 4; Supplemental Figure S6). To enable better penetration of DAPI, we also fixed and cleared the translational reporters before staining and imaging (Supplemental Figure S9), which confirmed the cytosolic location of all ACOs, and a nuclear localization for ACO1, 4 and 5 in roots.

**Figure 4:**
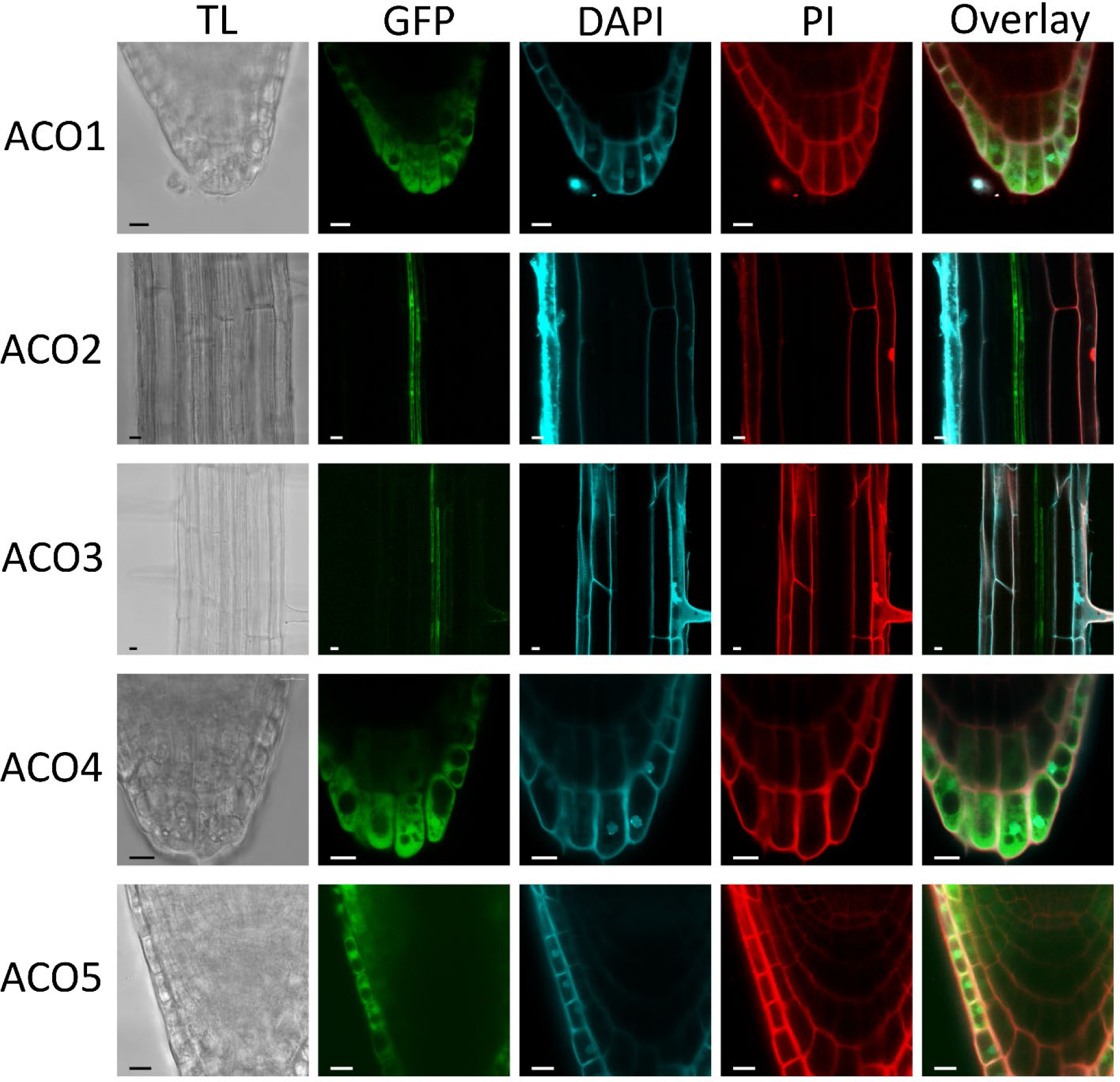
ACO proteins are subcellular localized in the cytosol and nuclei of roots. Confocal imaging of live 7-day-old ACO translational reporter lines (p*ACO1-5*:ACO1-5:GUS-GFP) after DAPI (nuclei) and PI (cell wall) staining. Rows show transmitted light (TL), GFP, DAPI, PI and an overlay. Scale bars are 10 μm.

### Ethylene has a limited feed-back on tissue-specific *ACO* expression patterns

We also investigated which ACOs are regulated by ethylene and ACC, because some *ACO* genes contain EIN3-binding sites in their promoter region. *ACO1*, *ACO4* and *ACO5* all harbor EIN3 binding sites within 1 kB upstream of the start codon, whereas *ACO2* and *ACO3* have EIN3 binding sites around 5.5 and 3 kB upstream of the start codon, respectively (Supplemental Figure S10). We therefore studied the tissue-specific expression in roots of dark-grown seedlings after a treatment with 1 ppm ethylene or 1 and 5 µM ACC for 4 h (Figure 5). Surprisingly, we did not see a lot of changes in expression levels of the *ACOs*, except for a slight increase of *ACO5* expression in the roots of dark-grown seedlings, indicating that *ACO5* is probably the only *ACO* gene that is substantially regulated by ethylene in dark-grown root tips. This is also supported by ChIP data which shows that EIN3 targets several regions of the *ACO5* promoter after ethylene treatment, whereas the other *ACO* promoters show fewer and smaller EIN3-binding ChIP peaks in seedling tissue (Supplemental Figure S10).

**Figure 5:**
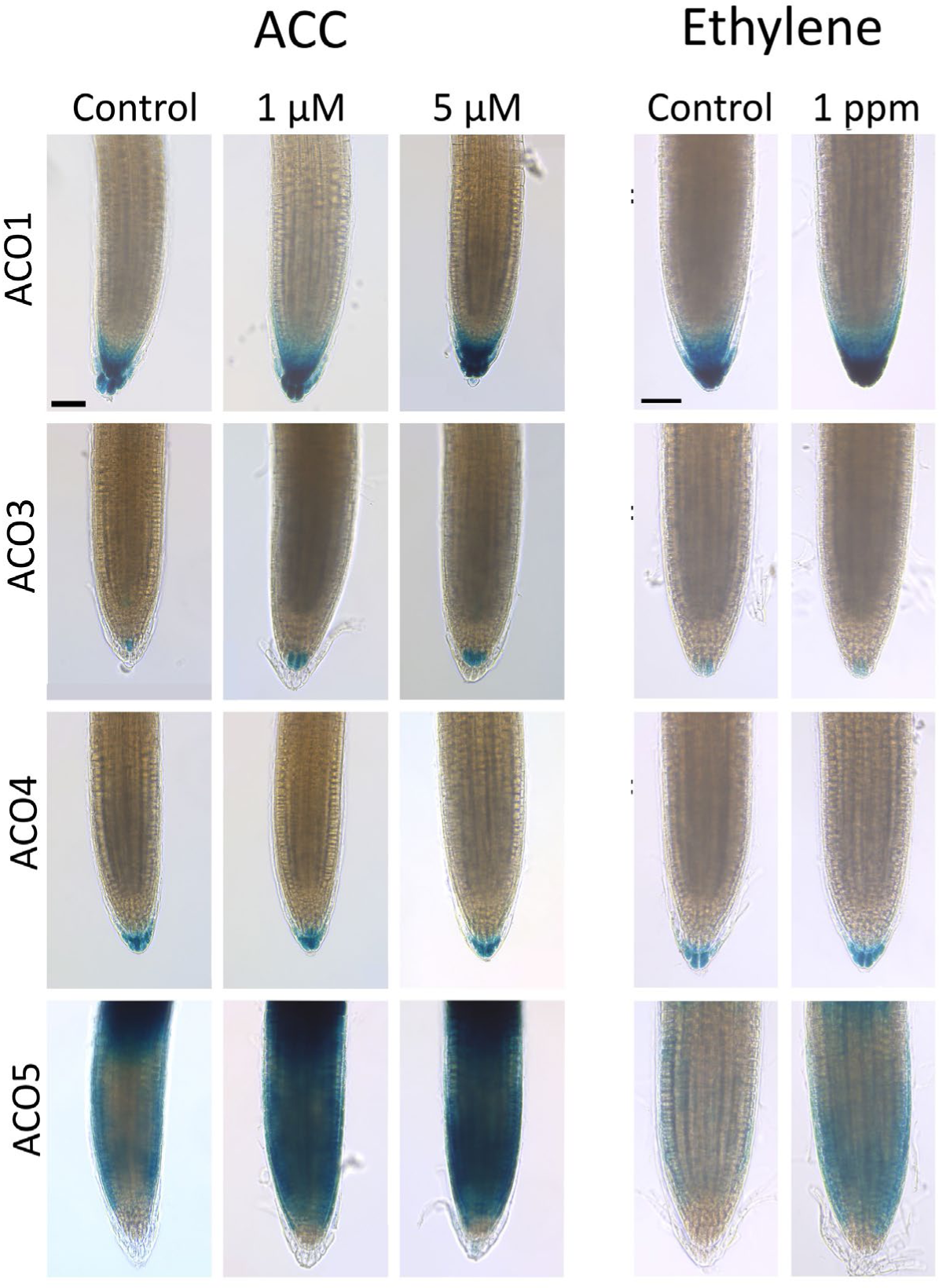
*ACOs* are not strongly transcriptionally regulated by ethylene or ACC in root tip. The *ACO* transcriptional reporter lines (p*ACO1-5*:*GUS-GFP*) were grown for 5 days in the dark and incubated with 1 ppm ethylene, or 1 or 5 µM ACC, for 4 hours before GUS staining. Scale bars are 50 µM.

### *ACO* gene redundancy dominates dark-and light-grown seedling development

To characterize the role of ACOs in the development of dark-grown seedlings, we performed a triple response assay. In a preliminary dose-response experiment, we found 2 µM ACC to be saturating for both root and hypocotyl development of 4-day-old dark-grown seedlings (Supplemental Figure S11). Therefore, we choose 0.2 µM as intermediate and 2 µM as a saturating concentration for testing all the *aco* single and higher order mutants (Figure 6). Most of the *aco* mutants did not show any strong hypocotyl phenotype compared to the wild type in the absence and presence of ACC, except for the quintuple lines (Figure 6A). With respect to the dark-grown roots, only the quintuple lines showed insensitivity towards a low and high dose of ACC (Figure 6B-C), indicating that a high level of *ACO* gene redundancy is at play during dark-grown hypocotyl and root development.

**Figure 6:**
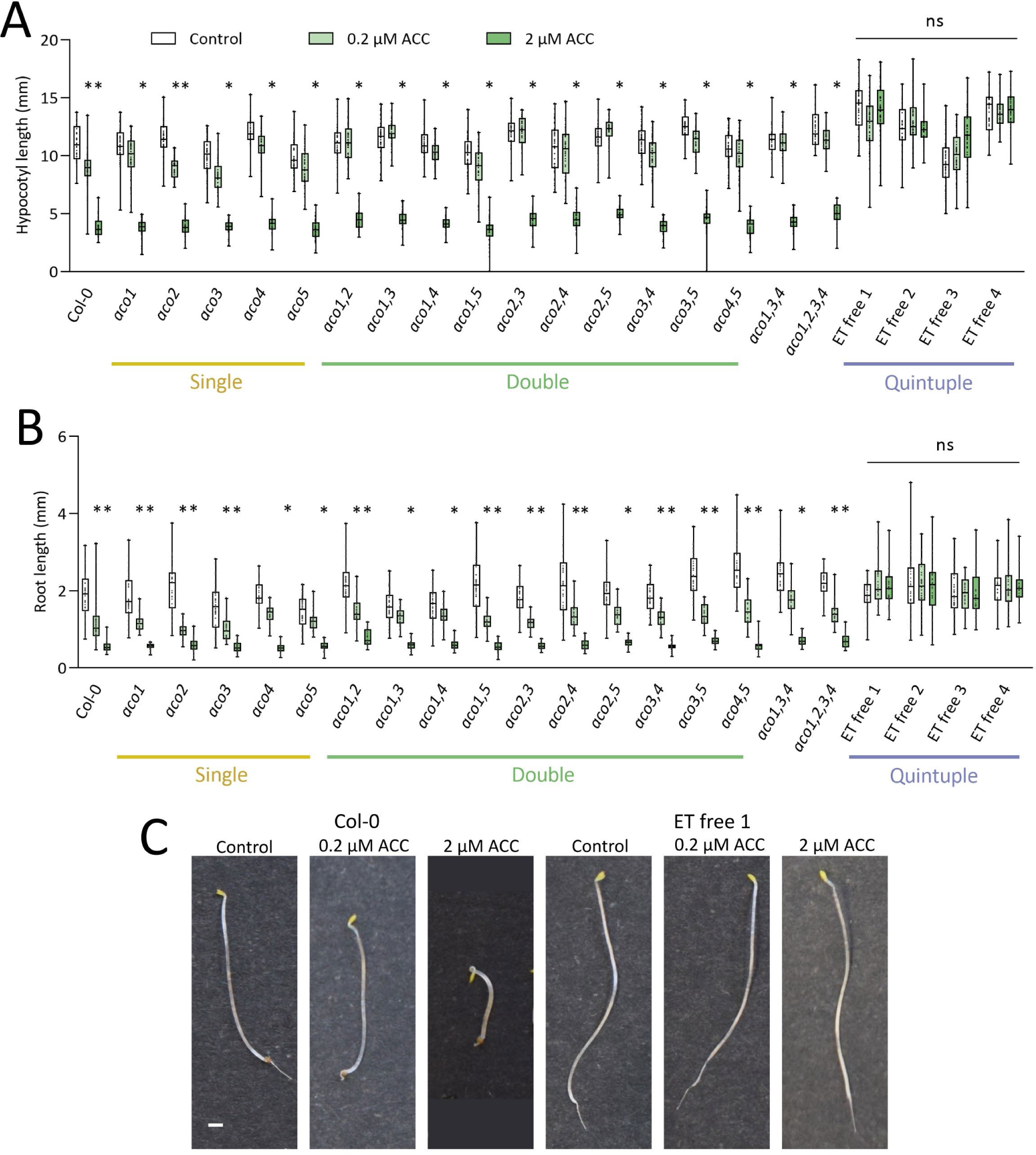
*ACO* gene redundancy during the triple response of dark-grown seedlings. Hypocotyl (A) and root (B) lengths of the wild type and single and higher order 4-day-old dark-grown *aco* mutants grown on different levels of ACC (0, 0.2 and 2 µM). (C) Representative seedlings of dark-grown wild type (Col-0) and *aco* quintuple (ET free 1) mutants at different ACC concentrations. Kruskal-Wallis and Dunn’s post-hoc test were performed per genotype (n ≥ 20). Asterisks indicate significant differences between the controls and ACC treatment per genotype (p < 0.05).

Next, we studied the genetic functionality of the *ACO* gene family during light-grown seedling development. Our preliminary dose response experiment (Supplemental Figure S11) showed that for 10-day-old light-grown seedlings 0.5 and 5 µM ACC were representative of an intermediate and saturating condition (Figure 7). Surprisingly, we did not observe a strong elongation of the hypocotyl of light-grown seedlings for the *aco* mutants except for *aco1,4* (Figure 7A), indicative that less ethylene was produced in the hypocotyl of *aco* mutants. For light-grown roots, we also did not observe striking differences between the wild type and the *aco* single and double mutants (Figure 7B). However, the quintuple *aco* mutant roots were insensitive to a low level of ACC (0.5 µM) and did not show a reduction in root length, indicative of their inability to produce ethylene. Interestingly, these *aco* quintuple lines did become sensitive again towards high levels of ACC (5 µM), as the roots of all ET free lines were small, similar as the wild type (Figure 7B-C). This suggests that ACC itself, when administered at high concentrations, can cause an inhibition of root elongation independent of ethylene, as the *aco* quintuple mutants are unable to convert ACC into ethylene. Altogether, we can conclude that a reduced or lack of ethylene biosynthesis does not drastically impact Arabidopsis seedling development.

**Figure 7:**
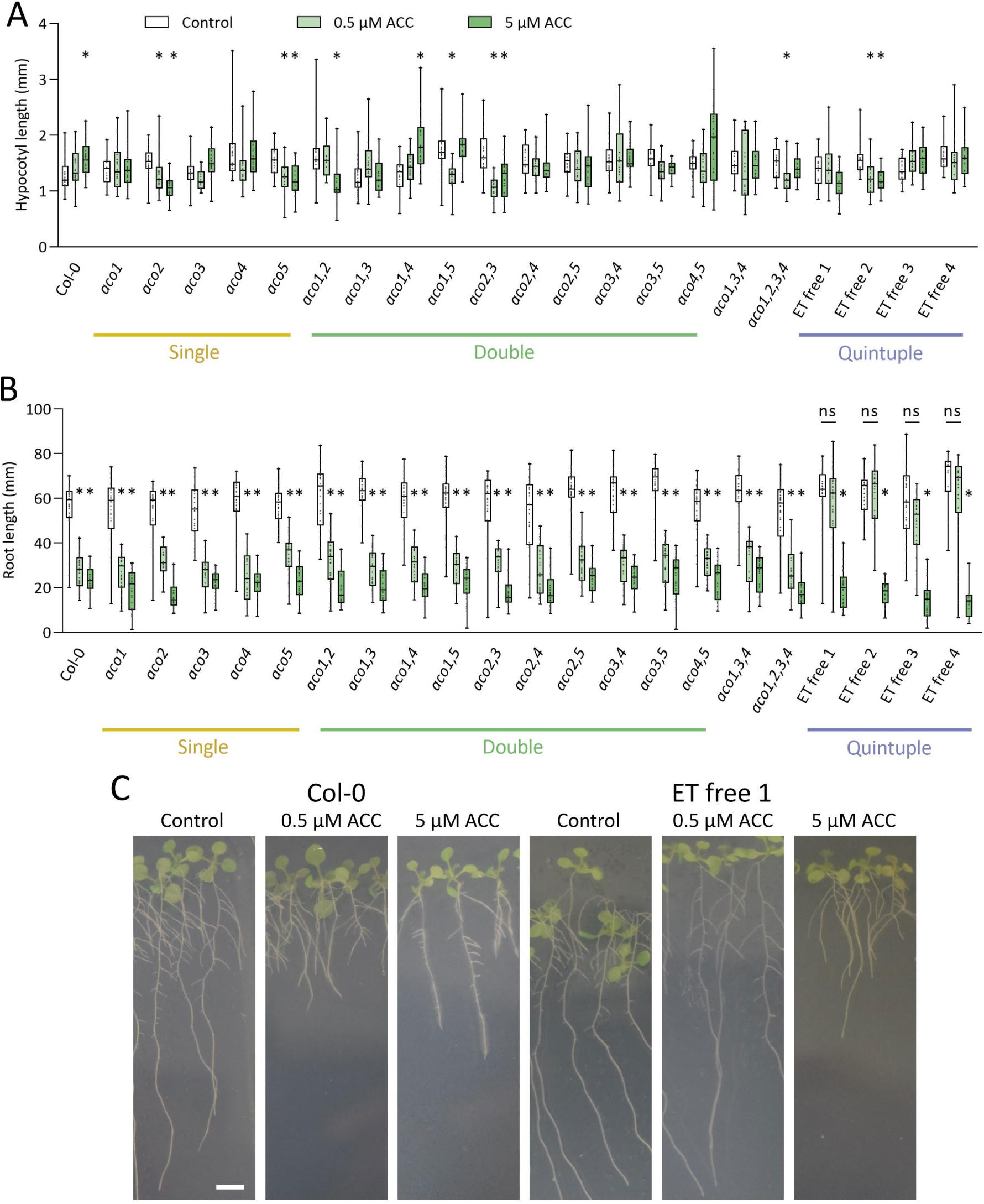
*ACO* gene redundancy during light-grown seedling development. Hypocotyl (A) and root (B) lengths of the wild type and single and higher order 10-day-old light-grown *aco* mutants grown on different levels of ACC (0, 0.5 and 5 µM). (C) Representative seedlings of light-grown wild type (Col-0) and *aco* quintuple (ET free 1) mutants at different ACC concentrations. Kruskal-Wallis and Dunn’s post-hoc test were performed per genotype (n ≥ 20). Asterisks indicate significant differences between the controls and ACC treatment per genotype (p < 0.05).

### Ethylene biosynthesis does not define vegetative and generative development

Besides the establishment of dark-and light-grown seedlings, we investigated the role of ACO during vegetative rosette development. We observed that some double mutants (*aco2aco3*, *aco2aco5* and *aco3aco4)* had a slightly larger rosette compared to wild-type plants (Figure 8A). However, the rosette of the *aco* quintuple lines was not consistently larger (Figure 8A; Supplemental Figure S12).

**Figure 8:**
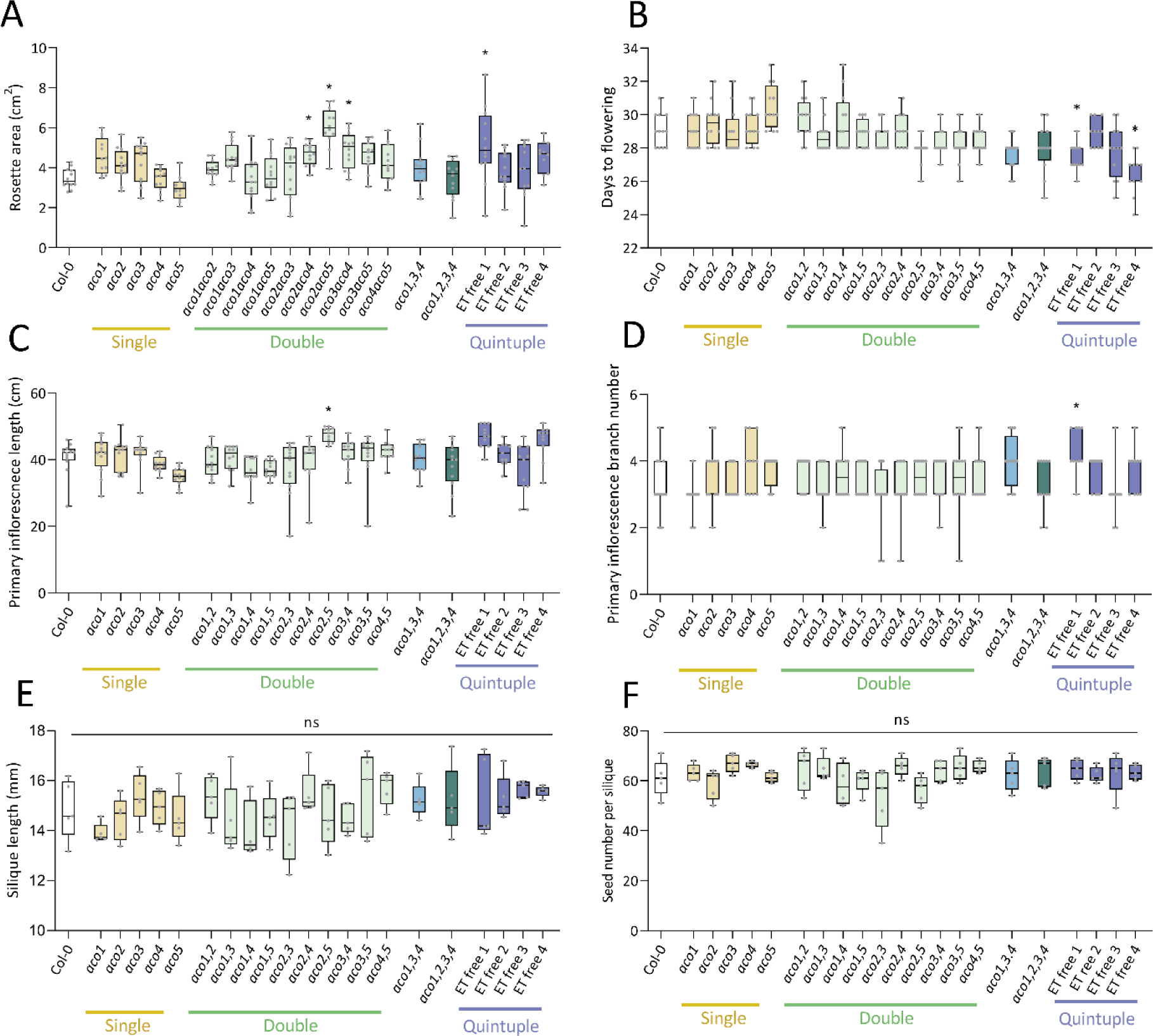
Vegetative and generative development of the *aco* mutants. (A) rosette size of 4-week-old plants, (B) days to flowering, (C) primary inflorescence length, (D) primary inflorescece branch number, (E) silique length, and (F) seed number per silique. Kruskal-Wallis and Dunn’s post-hoc test were performed per genotype (4 < n ≥ 20). Asterisks indicate significant differences between the wild type and the *aco* mutants (p < 0.05).

We also analyzed generative inflorescence and flower development and observed only very modest changes. Some ET free lines flowered one to two days earlier (Figure 8B) compared to wild-type plants. The primary inflorescence length (Figure 8C) and branching number (Figure 8D) did not change much between the different genotypes. The morphology of the ET free flowers was also identical to the wild type. The *aco* mutants developed normal siliques (Figure 8E) and had a normal seed-set number (Figure 8F). Altogether, we can conclude that a reduced or lack of ethylene biosynthesis does not impact vegetative and generative development of Arabidopsis.

### A lack of ethylene production delays but does not inhibit petal abscission

Besides vegetative and generative development, ethylene is often associated with senescence and abscission (Iqbal et al., 2017; and references therein), processes linked with high rates of ethylene production. Therefore we studied flower petal absission and found that the ET free lines abscised their petals on average 2 flower stages later compared to the wild type (Figure 9A-B). These results were similar to ethylene instensitive lines (*etr1-1* and *ein2-5*), which are known to have delayed petal abscission (Patterson & Bleecker, 2004). When studying the role of ethylene biosynthesis in rosette leaf senescence, we did not observe differences in the timing or rate of senescence between the wild type and two ET free lines (Figue 9C-D). These results were unexpected, but ethylene insentive mutants (*etr1-1* and *ein2-5*) also did not show any delay in leaf senescence.

**Figure 9:**
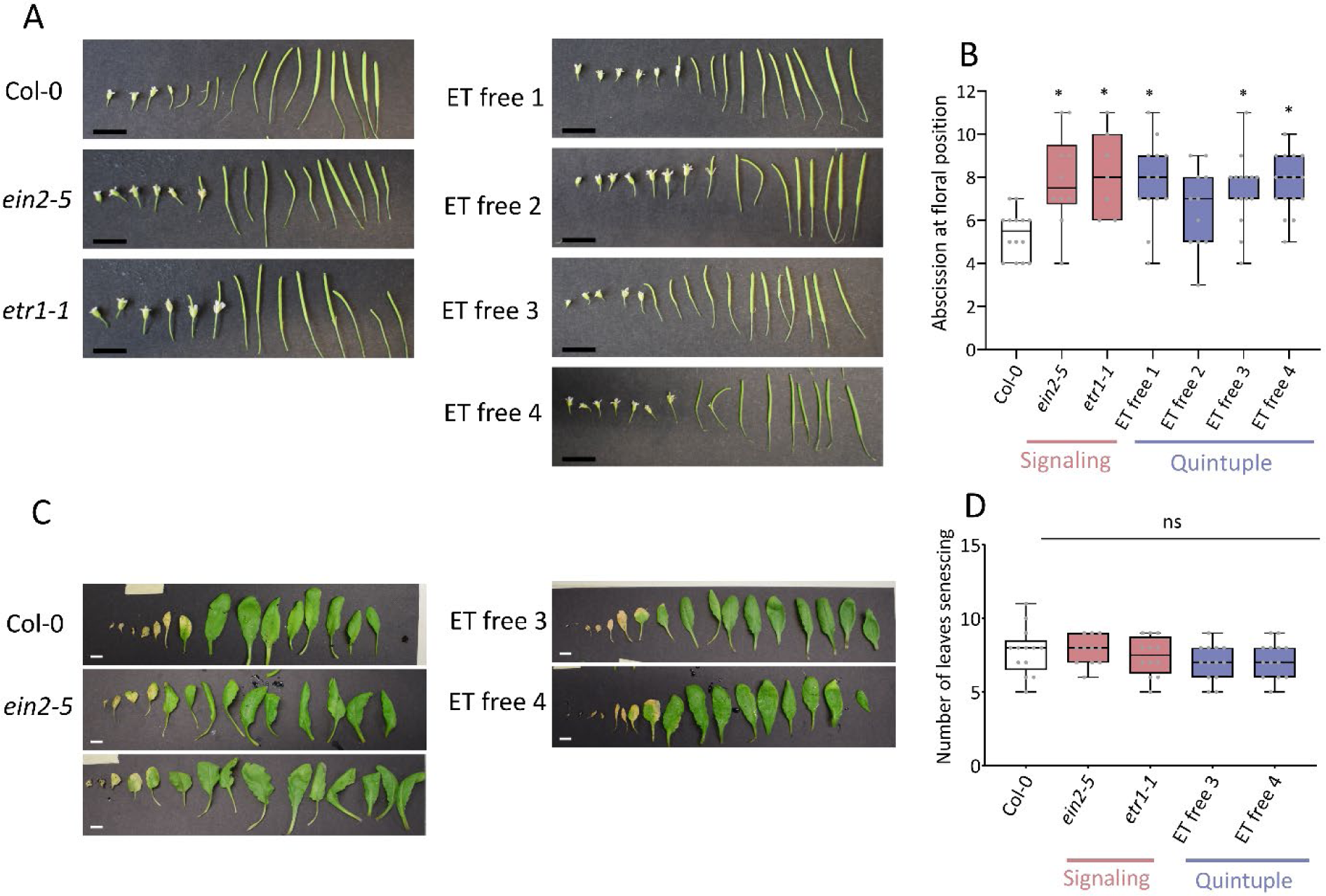
A lack of ethylene delays floral organ abscission, but shows no major effect on rosette leaf senescence. (A) Representative images of siliques at flower stages 1 – 16 and (B) the floral position of abscission (n ≥ 7). (C) Representative images of rosette leaves and (D) number of senescing leaves of 52-day-old plants (n ≥ 11). Scale bars in (A) are 10 mm and in (C) are 1 cm. Kruskal-Wallis and Dunn’s post-hoc test were performed per treatment, and asterisks indicate significant differences between the wild type and the *aco* mutants (p < 0.05).

### A lack of ethylene production impacts biotic and abiotic stress resilience

Previously, it was shown that ethylene signaling mutants perform worse during different biotic (Broekaert et al., 2006) and abiotic (Chen et al., 2022) stresses compared to wild-type plants, indicating that ethylene signaling plays an important role during stress resilience. Given our observations that a reduced or lack of ethylene production does not seem to alter normal vegetative and reproductive development of Arabiopsis, we wondered if ethylene synthesis might impact biotic and abiotic stress tolerance.

Our stress experiments showed that an inability to produce ethylene does no always alter biotic and abiotic stress tolerance for specific stressors. For salt stress (mild and severe), ET free lines did not show a great difference in root length compared to wild-type plants, while ethylene signaling mutants (*etr1-1* and *ein2-5*) did suffer more from salinity (Figure 10A). However, for nutrient deficiency stress, all ET free lines performed worse compared to the wild-type plants, similar to ethylene signaling mutants (Figure 10B). For heavy metal stress such as Al (Figure 10C; 100 µM) and Cd (Figure 10D; 20 µM) toxicity, the ET free lines performed better under mild stress, compared to wild-type plants, similar to the ethylene signaling mutants. When challenged with a biotic stressor, in this case a *Botrytis cinerea* leaf infection, ET free lines suffered more from the infection, similar to ethylene mutants, compared to wild-type plants (Figure 10E).

**Figure 10:**
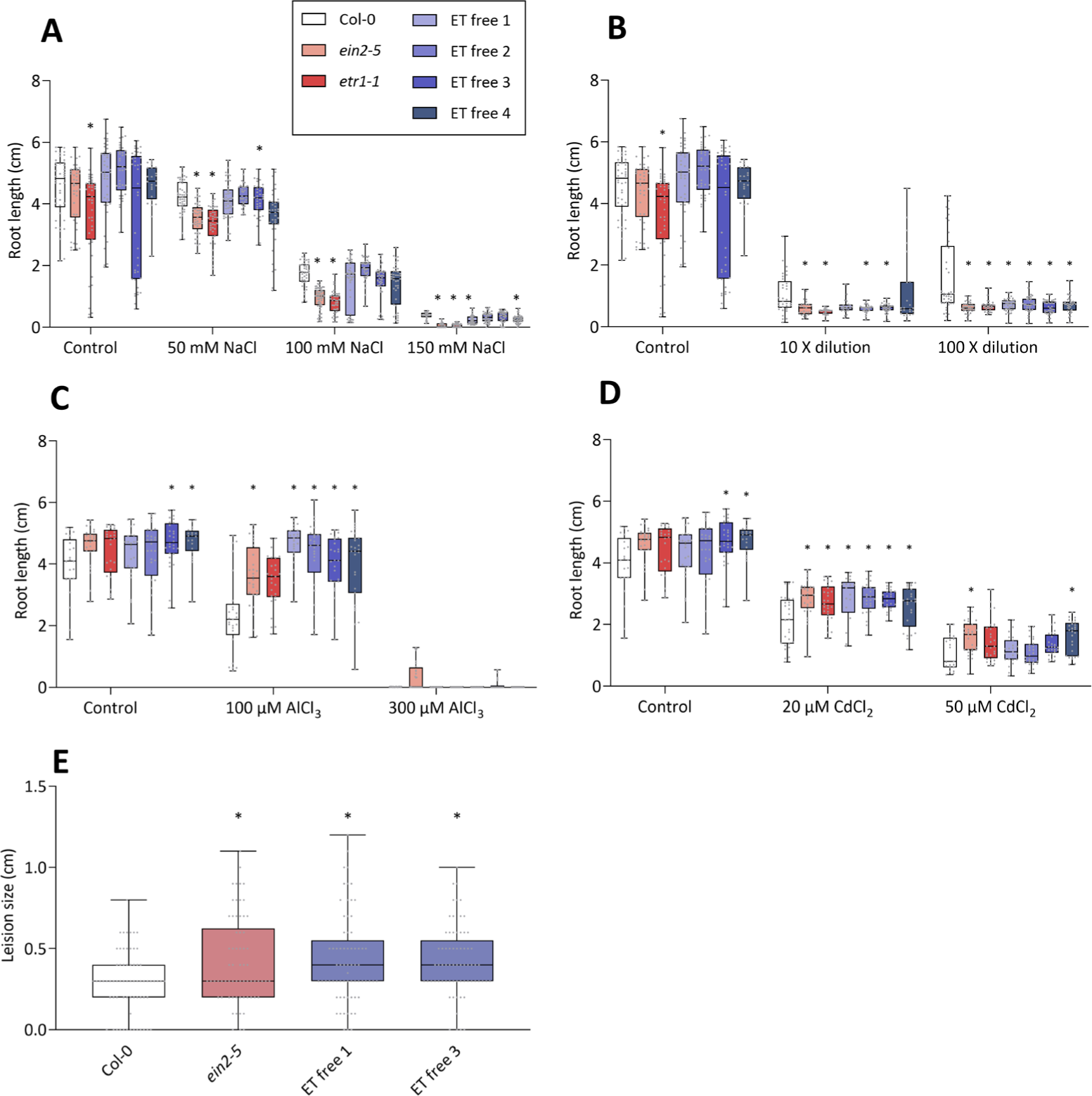
Loss of ethylene biosynthesis impairs biotic and abiotic stress resilience. Effect of different doses of (A) salt stress, (B) nutrient deficiency stress, (C) Al toxicity stress, and (D) Cd toxicity stress on 10-day-old seedling root development, and (E) biotic stress by means of a leaf infection with *Botrytis cinerae* (lesion size), for the *aco* quintuple mutants (ET free 1-4) and ethylene signaling mutants (*ein2-5* and *etr1-1*). Kruskal-Wallis and Dunn’s post-hoc test were performed per treatment (n ≥ 24) for A-D, and a Brown-Forsythe and Welch ANOVA analysis with a Games-Howell’s Multiple Comparisons post-hoc test was done for E (n ≥ 50). Asterisks indicate significant differences between the wild type and the *aco* mutants per treatment (p < 0.05).

## Discussion

### *ACO* gene redundancy sustains ethylene biosynthesis, and a lack of ethylene does not impair normal vegetative and reproductive development

For a long time, it has been assumed that ACS is the rate limiting step in ethylene biosynthesis (Yang and Hoffman, 1984), based on the observation that feeding external ACC leads to higher ethylene production rates. However, newer studies have shown that ACO can also be rate limiting during specific processes (e.g., during post-climacteric fruit ripening; Van de Poel et al., 2012). We now show that the *ACO* gene family has a very high level of redundancy amongst its members, and that ethylene levels are sustained in single, double, and higher order *aco* knock-out mutants. In fact, only when all five *ACO* genes are knocked-out, ethylene production levels are drastically impaired (Figure 1; Li *et al*., 2021). The insignificant or very small differences in ethylene production levels of single and higher order *aco* mutants align well with a lack of strong developmental phenotypes (seedling, vegetative and generative development). Even ET free lines, unable to produce ethylene, did not show strong developmental phenotypes (Figure 5-7). Only when the *aco* quintuple mutants were treated with external ACC, typical ethylene insensitive phenotypes were observed (e.g., no triple response). These data suggest that a lack of ethylene is not crucial for normal plant development, which makes us believe that ethylene is more a modulator of vegetative and reproductive development, in collaboration with other hormones through crosstalk (Van de Poel et al., 2015). Our results are in line with developmental data of several ethylene signaling mutants insensitive to ethylene (e.g., *etr1-1, ein2-5*, *ein3eil1*). These ethylene insensitive mutants also show a rather normal vegetative and reproductive development, with *etr1-1* having the most pronounced phenotypes (Alonso et al., 1999; Chao et al., 1997; Ogawara et al., 2003). Contradictory, constitutive ethylene signaling mutants (e.g., *ctr1-1*) or ethylene overproducing mutants (e.g., *eto1eol1eol2*) do show more drastic developmental phenotypes (Kieber et al., 1993; Achard et al., 2007; Christians et al., 2009; Ogawara et al., 2003). These observations suggest that a lack of ethylene production or sensitivity do not impair normal vegetative development, while enhanced ethylene production or sensitivity do.

Interestingly, we observed differences in petal abscission but not in leaf senescence, two processes well-known to be associated with high rates of ethylene production (Grbic & Bleecker, 1995; Patterson & Bleecker, 2004). Although leaf senescence is known to be associated with ethylene sensitivity (Grbic & Bleecker, 1995), our experiments contradict this finding, and more research is needed to explain this. However, we conclude that a lack of basal levels of endogenous ethylene do not impair normal plant development and reproduction, but only some processes where high rates of ethylene are produced (e.g., petal abscission), or triggered by external ACC feeding, rely on a fully functional ACO protein family. These observations suggest that ACS and the production of ACC is likely key to direct basal ethylene levels and hence normal plant development, but that both ACS and ACO need to cooperate to activate specific developmental processes such as abscission, during which high rates of ethylene are produced. The role of ACOs in other processes associated with high rates of ethylene synthesis in different plant species, such as climacteric fruit ripening (Alexander & Grierson, 2002) remain to be studied. Knocking out one ACO (*SlACO1*) in tomato is enough to confer 95 % of ethylene synthesis and drastically delay fruit ripening (Hamilton et al., 1990), indicative that ACOs play a prime role in ethylene synthesis during climacteric fruit ripening.

### Ethylene production is tissue-specific and developmentally programmed

Ethylene, as a modulator of plant growth and development, should be produced in a coordinated way to drive tissue-specific responses. The tissue specificity of ethylene production was previously assessed biochemically by measuring levels of ethylene, ACC and its conjugates of dissected tissues of tomato fruit (Martin *et al*., 1995; Van de Poel *et al*., 2014*b*). Our work now contributes to this spatial aspect by revealing the developmental-specific and tissue-specific expression and localization of the different *ACO* genes in Arabidopsis (Figure 3, 5). Comparing our results with previous studies about the tissue-specific expression of the *ACS* gene family (Tsuchisaka & Theologis, 2004; Wang et al., 2005) reveal that for certain tissues (such as root tips, root vasculature, root-shoot junction, hypocotyl, leaf mesophyll, anther filaments, and stigma) *ACS* and *ACO* expression align spatially and developmentally. This indicates that the full and dedicated ethylene biosynthesis pathway (ACS + ACO) are co-localized and suggests that ethylene biosynthesis is directed to specific developmental stages and tissues.

To produce ethylene, a gene needs to be translated into an active enzyme. Our work reveals that different ACOs have different affinities towards their substrate ACC (different K_m_ values; Figure 2) and that not all enzymes are equally active (different V_max_ values; Figure 2). For the ACS family, different members also have different K_m_ values ranging between 8-45 µM (Yamagami et al., 2003). Therefore, we can conclude that, besides a tissue-specific gene expression, isoform-specific ACO and ACS activity contribute to the precisely control of actual ethylene production rates. For example, ACO1 is the most active ACO isoform with a rather high affinity for ACC (K_m_ = 33 µM) and is highly expressed in the root tip (of both light-and dark-grown seedlings). Interestingly, only *ACS8* is expressed in root tips (Tsuchisaka & Theologis, 2004), but ACS8 also has a rather high affinity (K_m_ = 15 µM). These results might indicate that the root tip tissue has a potential very high and localized ethylene emission rate. Alternatively, ACO5 has a much lower ACC affinity and a lower ethylene production capacity (Figure 2) but has a broad expression pattern in the root. Therefore, ACO5 might be an isoform that contributes to a steady but low level of basal ethylene production.

Interestingly, ethylene and ACC do not exert a major feedback effect on the tissue-specific *ACO* expression in roots (Figure 5), suggesting that the *ACO* gene family of Arabidopsis is likely regulated by other hormones and not ethylene itself. In fact, it is well established that ethylene crosstalks with auxins to regulate primary root development (Stepanova et al., 2007; Swarup et al., 2007; Ruzicka et al., 2007), although the role of ACOs in this remains unstudied. Furthermore, cytokinins activate tissue-specific expression of *ACO2*, *ACO3* and *ACO4* in Arabidopsis roots leading to an inhibition of primary root growth (Yamoune *et al*., 2023). Brassinosteroids activate *ACO1* expression in the root tip via BES1, and contribute to the regulation of a gravitropic response (Park *et al*., 2020). Oppositely, brassinosteroids inhibit ethylene production in the shoots by down regulating *ACO2* and *ACO4* expression directly via the transcription factor BES1 and BZR1 respectively (Moon *et al*., 2020; Moon et al., 2021). These examples show that ACO is tightly regulated by other hormones and indicate that ACO is a key player in establishing tissue-specific ethylene synthesis that allows fine-tuning of local developmental and growth responses through hormonal crosstalk.

### Ethylene is likely produced in the cytosol, but ACOs also reside in the nucleus

Despite the discovery decades ago that plants produce ethylene (Gane, 1934), the actual sub-cellular localization of ethylene synthesis in plants remains contested. Some early studies using ACO activity assays, linked the production of ethylene with the vacuole and specifically the tonoplast (Guy and Kende, 1984; Mayne and Kende, 1986). In those days the purification of ACO was troublesome and ACO enzyme activity was thought to be dependent on membrane integrity, hence suggesting that ethylene production was associated with membranes (Kende, 1989). Experiments with radiolabeled ACC showed that ethylene synthesis could either be cell membrane-associated or cytosol-associated, depending on the plant species (Bouzayen *et al*., 1990). Soon thereafter, it was discovered that the supplementation of vitamin C to the ACO extraction protocol was crucial to retain enzyme activity (Ververidis and John, 1991). Perhaps enzyme preparations of membrane fractions lost ACO activity due to a lack of an essential cofactor. However, immunolocalization studies using anti-ACO antibodies in apple and tomato cells and protoplasts, supported the belief that ACO is membrane-associated and likely apoplastically localized (Rombaldi *et al*., 1994; Ramassamy *et al*., 1998). Contradictory studies showed that ACO, and hence ethylene production, was localized to the cytosol. Using other anti-ACO antibodies, the subcellular localization of ACO was confined to the cytosol of tomato suspension cells (Reinhardt *et al*., 1994), apple fruit (Chung et al., 2002), and Douglas fir stems (Hudgins et al., 2006). The discovery of the first *ACO* gene also suggested it codes for a cytosolic protein (Hamilton et al., 1990). Now, our results, using stable translational reporter lines in Arabidopsis, in which ACO was fused to a GFP protein and driven by its native promoter, revealed that Arabidopsis ACOs are localized to the cytosol and nucleus of root cells (Figure 4; Supplemental Figure S9). This supports in part a previous study that ectopically overexpressed (*35S*) the safflower *SmACO1:GFP* in onion cells and found that ACO localizes primarily to the cytosol (Tu *et al*., 2019), but they also retrieved a strong nuclear SmACO1:GFP signal in some cells, matching our data. These results are surprising, as ACOs do not have any known nuclear localization peptide, and hence their nuclear location is established by an unknown mechanism. It also remains to be studied if ACS is also nuclear localized or if ACC itself is shuttled into the nucleus to enable nuclear ethylene production. And if so, the role of nuclear ethylene synthesis remains mysterious. Alternatively, ACO might have another moonlight function in the nucleus besides ethylene synthesis.

Nonetheless, our data in combination with previous histological immunolocalization studies, provides compelling evidence that ethylene is produced in the cytosol. In fact, 2-oxoglutarate-dependent dioxygenase, the larger enzyme family to which ACO belongs, are typically cytosolic enzymes (Kawai *et al*., 2014).

### The *aco* quintuple mutant reveals an ACC-specific phenotype in primary root development

Lately, attention is given to the role of ACC as a signaling molecule independent of ethylene (Polko and Kieber, 2019; Li *et al*., 2022*b*). This novel role for ACC was brought to light when more plant genomes were sequenced and revealed that non-seed plants lack homologs of *ACO* (Li et al., 2022*b*), questioning their capacity to convert ACC into ethylene. Interestingly, non-seed plants are capable of producing ACC and contain ACS homologs, but they produce ethylene via a different pathway, indicating that ACC has a function outside ethylene biosynthesis (Van de Poel, 2020; Li *et al*., 2022*b*). This ACC independent function has been revealed in several plant species including a red algae (*Pyropia yezoensis*; Uji *et al*., 2020), a liverwort (*Marchantia polymorpha*; Li *et al*., 2020; Katayose et al., 2021), tomato (Althiab-almasaud *et al*., 2021), and Arabidopsis (Xu *et al*., 2008; Tsang *et al*., 2011; Vanderstraeten *et al*., 2019; Yin *et al*., 2019; Mou *et al*., 2020). In the latter species, ACC controls root development (Xu *et al*., 2008; Tsang *et al*., 2011), stomatal development (Yin *et al*., 2019), pollen tube attraction (Mou *et al*., 2020) and vegetative development (Vanderstraeten *et al*., 2019.) We now revealed that ET free Arabidopsis plants, incapable of producing ethylene due to the loss of all 5 *ACO* genes, have ACC-specific responses. We observed that light-grown seedlings were sensitive to high levels (5 µM) of ACC, leading to a strong inhibition of root development independent of ethylene synthesis, suggesting that ACC itself caused this phenotype. Our results corroborate previous discoveries where ACC impairs primary root development by using ACO inhibitors and/or inhibitors of ethylene signaling and ethylene insensitive mutants (Xu *et al*., 2008; Tsang *et al*., 2011; Vanderstraeten *et al*., 2019). It remains unclear how changes in ACC levels impact primary root development, but some pioneering studies have suggested that ACC probably acts by sensing/influencing cell wall polymer composition/flexibility (Xu *et al*., 2008; Tsang *et al*., 2011).

### Ethylene production is needed to facilitate biotic and abiotic stress resilience

Ethylene is well known as a stress hormone, for both biotic and abiotic stressors (Broekaert et al., 2006; Chen et al., 2022) and this role of ethylene in stress responses is evolutionary conserved in plants (Van de Poel & de Vries., 2023). It is well established that ethylene signaling mutants, such as ethylene insensitive (e.g., *etr1-1*, *ein2-5*, and *ein3eil1*) and constitutive ethylene sensitive (e.g., *ctr1-1*) mutants have an altered stress resilience against salinity (Riyazuddin et al., 2020), flooding (Sasidharan & Voesenek, 2015), metal toxicity (Keunen et al., 2016), nutrient deficiency (Garcia et al., 2015) and temperature (Huang et al., 2023), as well as many biotrophic and necrotrophic pathogens (van Loon et al., 2006). This made us wonder if a lack of ethylene biosynthesis also plays a prominent role in mediating biotic and abiotic stress responses. Our results showed that a lack of ethylene synthesis does not impair salinity stress responses (Figure 10) but is involved in nutrient deficiency and heavy metal stress responses, as well as resistance towards the fungal pathogen *Botrytis cinerea* (Figure 10). In fact, many stress responses of ET free lines were similar to stress responses of ethylene insensitive lines (*etr1-1* and *ein2-5*), suggesting a similar role for both ethylene production and signaling in facilitating stress responses.

## Acknowledgement and funding

This work was financially supported by KU Leuven with a start-up grant (STGBF/16/005) and research grant (C14/18/056) to B.V.d.P., a postdoctoral fellowship to S.R. (PDMT2/22/035), as well as the Research Foundation Flanders with research grants to B.V.d.P. (G092419N; G0G0219N; G023124N), a PhD fellowship to M.H. (SB1S18717|SB1S18719), J.P. (1150822N|1150824N), and a postdoctoral fellowship to W.M. (1207022N) and S.R. (12AKQ24N).

## Data availability

All ACO reporter lines and *aco* mutant generated in this study are available through the Nottingham Arabidopsis Stock Center (NASC). All other data is available upon a reasonable request.

## Author contribution

MH, JVH, BDC, JJK, AV and BVdP designed the experiments, MH, JVH, JP, WM, SR, ARM, EK, SR and AP performed the experiments. JVH and BVdP wrote the manuscript.

## Supplementary Material

Supplemental Methods: Details on certain methods used in this study.

Supplemental Figure S1: Overview of the single *aco* mutants and the *ACO* gene expression. Supplemental Figure S2: Sites of CRISPR-induced mutations in *ACO5* in the ET free 3 and 4 lines. Supplemental Figure S3: Light-grown ET free lines do not produce ethylene.

Supplemental Figure S4: Coomassie stained SDS-PAGE gels of the recombinant ACO purification steps. Supplemental Figure S5: *ACO1* expression is confined to the close proximity of apical meristems.

Supplemental Figure S6: ACO2 translational reporter shows phloem-specific localization.

Supplemental Figure S7: Single-cell RNA-seq (scRNA-seq) data confirms *ACO* expression patterns observed in the *ACO* reporters.

Supplemental Figure S8: ACOs show differential expression in rosette leaves. Supplemental Figure S9: ACOs show cytosolic and nuclear subcellular localization. Supplemental Figure S10: The *ACO* promoters harbor EIN3 binding sites.

Supplemental Figure S11: Phenotyping of Col-0 wild type seedlings to an ACC gradient treatment.

Supplemental Figure S12: Representative images of rosettes of 3-and 4-week-old plants of Col-0 and the different *aco* quintuple lines (ET free 1-4).

Supplemental Table S1. Seed stocks used in this study. Supplemental Table S2. Primers used in this study.

## Notes

### Competing Interest Statement

The authors have declared no competing interest.

