## Supplementary Material for "1-Aminocyclopropane-1-carboxylic acid oxidase determines the fate of ethylene biosynthesis in a tissue-specific way to fine-tune development and stress resilience"

### Supplementary Methods:

#### Plant growth conditions

Before germination, seeds were surface sterilized for 2 minutes in 70 % ethanol (v/v) and 5 minutes in a 5 % bleach solution (NaOCl; w/v), followed by washing in sterile $H_2O$. Next, seeds were sown on Petri dishes containing sterile 0.5 x Murashige and Skoog (MS) basal salt medium with 1 % agar (w/v; pH 5.7), with or without different doses of ACC (Abcam), followed by imbibition at 4 °C for three days. For soil cultivation, 5-day-old plate-grown seedlings were transplanted to soil (Zaaien en stekken; DCM). Plants were grown under sunlight-mimicking LEDs (100 µmol.$m^{-2}$.$s^{-1}$) with a day/night cycle of 16 h/8 h at a constant temperature of 21 °C and 60-70 % relative humidity. For infection assays involving *Botrytis cinerea*, plants were grown under sunlight-mimicking LEDs (120 µmol.$m^{-2}$.$s^{-1}$) with a day/night cycle of 12 h/12 h, and with a day/night temperature of 23 °C/21 °C, and 60-70 % relative humidity.

#### ACO reporter line cloning and plant transformation

For the creation of the transcriptional *ACO* reporter lines, the promoter region (1807 - 2769 bases upstream from the ATG start codon) of each *ACO* gene was amplified.. Promoters were cloned into pDONR221 and then pKGWFS7 (carrying a GUS:GFP fusion) using Gateway cloning (primers see Supplemental Table S2). For the creation of the translational ACO reporter lines, the same promoter sequences as used for the transcriptional reporters were cloned into the BsaI restriction sites of the greengate (Lampropoulos *et al.*, 2013) vector *pGGA000*. The CDS (excluding stop codon) of the ACOs were cloned into the BsaI sites of *pGGC000*. Any native BsaI restriction sites within the native promoters or CDS were removed by site directed mutagenesis before cloning to the greengate vectors. The A and C module greengate vectors with promoters and CDS of the *ACOs* cloned were used for BsaI based greengate cloning to generate N- and C-terminal GFP translational reporters. All constructs were transformed into *Arabidopsis thaliana* using the *Agrobacterium* (strain GV3101) floral dip method. Transcriptional reporters were transformed into ecotype Col-0 and translational reporters transformed into their respective T-DNA mutants. Homozygous T3 lines were selected for use in experiments.

**RNA extraction and qPCR**

Briefly, RNA was extracted from 10-day old light-grown seedlings using the GeneJET RNA purification kit (Thermo Scientific) followed by a DNase treatment (RapidOut DNA Removal Kit, Thermo Scientific). Next, cDNA was made using the iScript™ Kit (Bio-Rad) according to the manufacturer's guidelines. RT-qPCR was performed using specific primer pairs (Supplemental Table S2) using the Green super mix (SsoAdvanced Universal SYBR™, Bio-Rad) on the CFX96 touch real-time PCR machine (Bio-Rad). The relative expression was quantified using the delta-delta Ct method, and the average expression of 4 replicates was normalized against the average expression of four reference genes (*UBQ10*, *UBC21, TIP41* and *YLS8*).

**GUS staining**

For GUS staining, plants were incubated in cold 90 % acetone (v/v) for at least 30 minutes on ice, then washed twice in staining buffer (50 mM $NaHPO_4$, 2 mM potassium ferricyanide, 2 mM potassium ferrocyanide, 0.2 % (v/v) Triton X-100; pH 7). Next, plants were incubated in staining buffer with 2 mM X-gluc at 37 °C in the dark until color development was observed. Following GUS staining, plants were cleared in 70 % ethanol (v/v) for at least 2 days. GUS-stained plants were imaged using a Leica Thunder Imager or a stereomicroscope (Olympus SZX).

**ACC and MACC quantification**

Briefly, ACC and MACC were extracted from 600-900 mg frozen crushed tissue using a double volume of ice-cold 80 % ethanol (v/v). The mixture was vortexed briefly and incubated for 1 h at 4 °C with gentle shaking, followed by 10 min centrifugation at 17000 x g. For ACC (500 µL) and for MACC (250 µL) supernatants were taken and vacuum evaporated overnight. For ACC, the pellet was dissolved in 500 µL water and was directly used for quantification. For MACC, the pellet was dissolved in 100 µL water and first hydrolyzed into ACC. This was done by loading the sample onto a 300 µL Dowex cation-exchange column (50WX8, 100-200 mesh) to remove residual ACC, followed by an acid hydrolysis with 80 µL 6 M HCl by boiling for 3 h at 100 ° C. Subsequently, the sample was neutralized by the addition of 80 µL 6 M NaOH. The ACC conversion into ethylene gas was done by incubating 1100 mL of sample (diluted with water) with 100 µL 5 % NaOCl-NaOH (2:1, v/v) diluted 50 % with water (v/v) and 100 µL $HgCl_2$ (20 mM) for 4 min at 4 °C. The sample was briefly vortexed before and after the incubation time, and 1 mL of headspace was sampled for ethylene quantification by GC as described above.

**Recombinant ACO protein purification**

E. coli cultures (2 L) expressing the recombinant ACOs after IPTG feeding, were pelleted by centrifugation at 5,500 rpm for 30 min at 4 °C, resuspended in lysis buffer, rotated at room temperature for 30 min, and sonicated with a Branson 450 sonifier (VWR) with a power of 3 and duty cycle of 80, for six cycles of 1 min sonication and 2 min rest on ice. Cell debris was removed by centrifugation for 40 min at 13,000 rpm at 4 °C and filtration through a 0.45 µm filter. The ACOs were first purified by IMAC chromatography (Qiagen Ni-NTA, 5–250 mM Imidiazole, 200 mM NaCl, 50 mM $NaH_2PO_4$, 1 mM DTT, pH 8) followed by anion exchange chromatography on an Äkta Prime Plus chromatography system (GE Healthcare) fitted with a 5 mL HiTrap Q HP column (Cytiva) (25–1000 mM NaCl, 50 mM $NaH_2PO_4$, 1 mM DTT, pH 8). SDS PAGE confirmed protein purity at every step. Protein concentration was calculated using a NanoDrop 2000 spectrophotometer (Thermo Scientific).

***Botrytis cinerea* sporulation**

*B. cinerea* B05.13 spores stored at -80 °C were plated on 0.5 x potato dextrose agar (PDA) plates (Difco™ Potato Dextrose Agar, BD). The plates were incubated in the dark at 25 °C for 7 days. Sporulation was induced by placing the *B. cinerea* plates under an UV-A lamp overnight at room temperature followed by incubation of the *B. cinerea* plates in the dark at 25 °C for 7 days. Spores were harvested by washing the *B. cinerea* plates using sterile $H_2O$ and a scraping off spores with a sterile spatula. The solution was filtered using sterile glass-wool and collected in 50 mL falcon tubes. The filtered spores were centrifuged for 20 minutes at 3970 x g at 20 °C. The supernatant was discarded followed by washing the spores twice in sterile $H_2O$. The spores were resuspended in 500-1000 µL of sterile $H_2O$ and spore concentration was evaluated using a Thoma-chamber. The spore solution was resuspended in an equal volume of 50 % glycerol solution (v/v), aliquoted in cryo-tubes and stored at -80 °C till use.

### Supplementary Figures

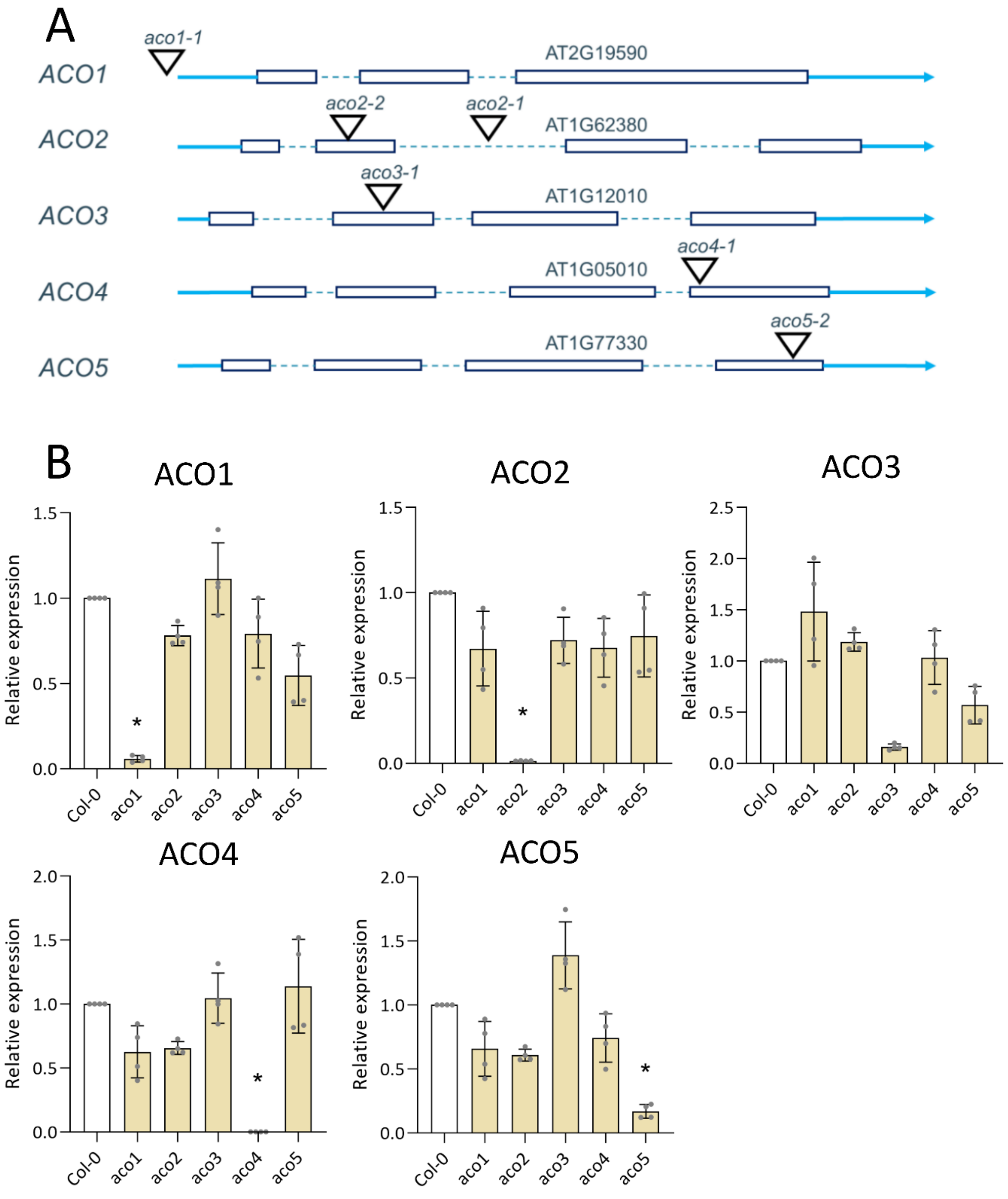


Supplemental Figure S1: Overview of the single *aco* mutants and the *ACO* gene expression. (A) Schematic representation of the *ACO1-5* gene models and the associated T-DNA positions. (B) Relative expression (qRT-PCRs) of the *ACO1-5* genes in the different *aco* single mutants of 10-day old light-grown seedlings (n = 4). Data is normalized against the average expression of 4 housekeeping genes (*UBC21, UBQ10, TIP41, YLS8*) per sample and analyzed with the delta-delta CT method. Significant differences in normalized expression between the wild type and the single *aco* mutants were tested using Kruskal-Wallis and Dunn's multiple comparisons test and are depicted by an asterisk ($p<0.05$).

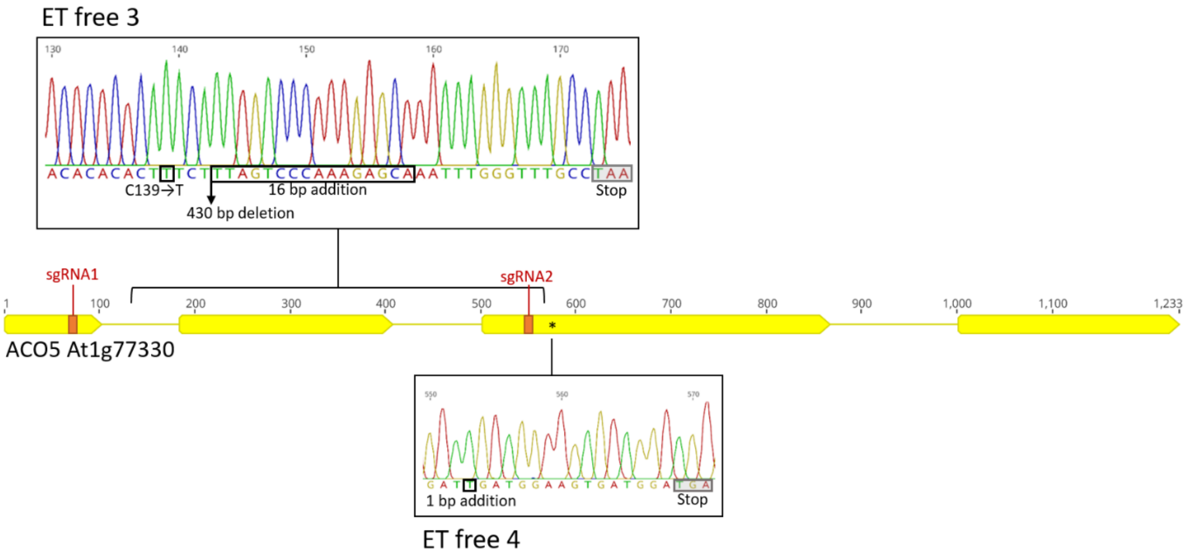


Supplemental Figure S2: Sites of CRISPR-induced mutations in *ACO5* in the ET free 3 and 4 lines. Mutations were identified by amplicon sequencing of a region of *ACO5* including the two sgRNA target sites (labelled in red). Boxed bases indicate mutation sites, and the premature stop codons generated are indicated in gray boxes.

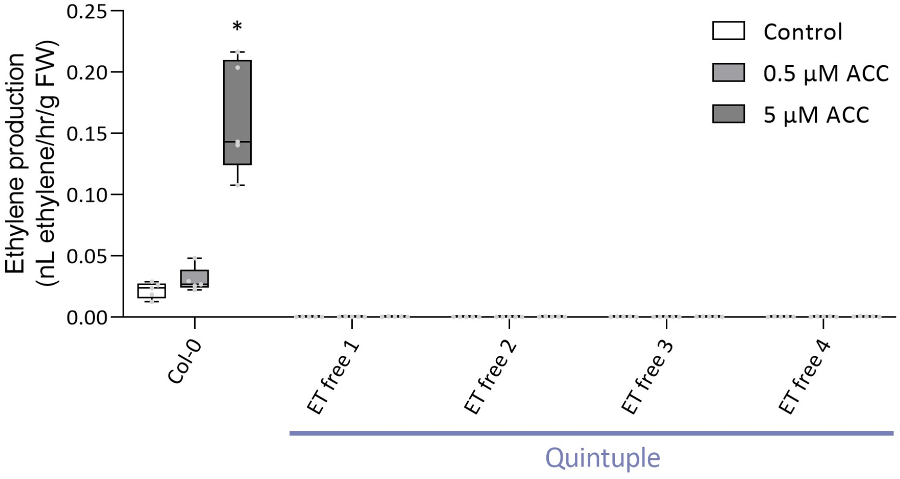


Supplemental Figure S3: Light-grown ET free lines do not produce ethylene. Ethylene production rates of 11-day-old light-grown wild type and *aco* quintuple mutant (ET free 1-4 seedlings) seedlings (n = 5) in the presence or absence of ACC (0.5 and 5 µM). Kruskal-Wallis and Dunn's post-hoc test were performed per genotype. Significant differences between the different treatments are depicted by an asterisk ($p<0.05$).

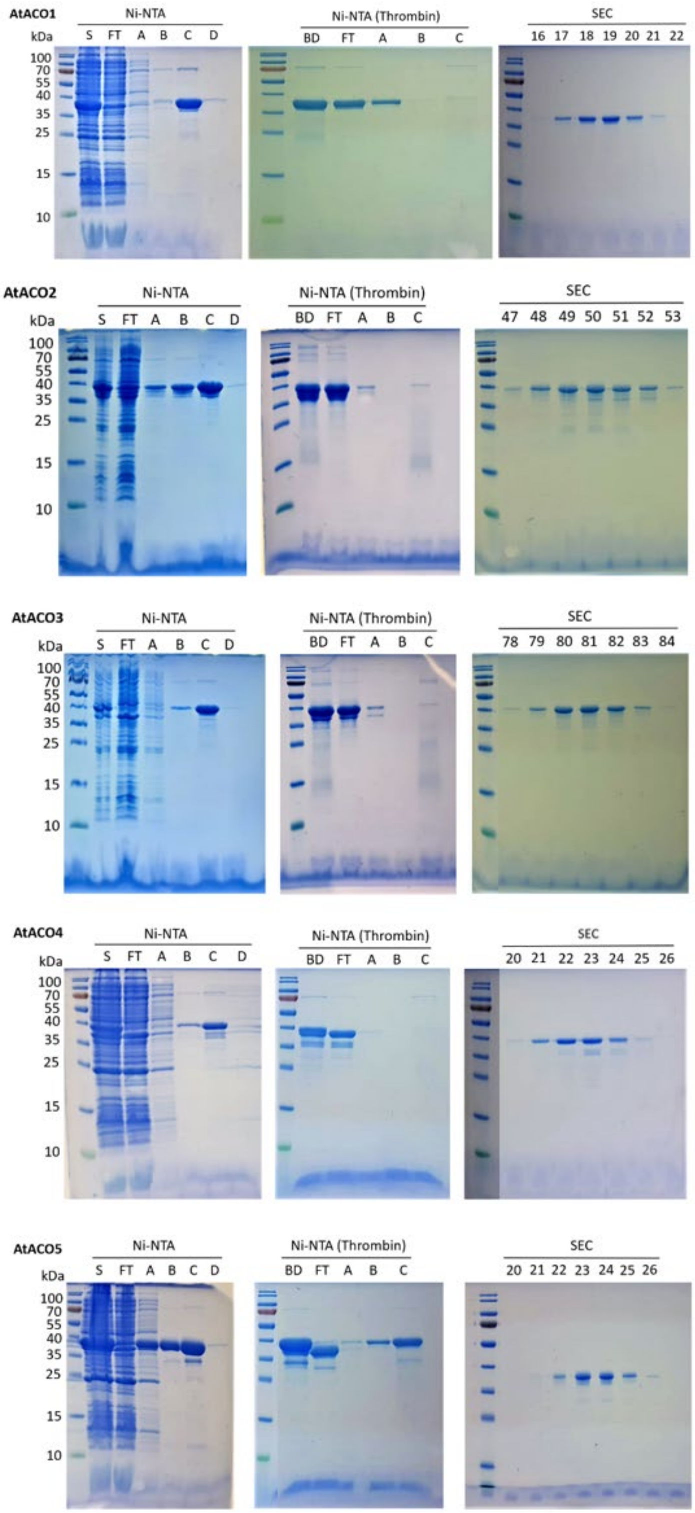


Supplemental Figure S4: Coomassie stained SDS-PAGE gels of the recombinant ACO purification steps. The gels of 3 different purification steps (Ni-NTA, Ni-NTA after thrombin cleavage and SEC (size exclusion chromatography) are shown. S: supernatant, FT: flow-through, A, B, C, D: protein elution fractions at increasing imidazole concentrations, BD: before dialysis. The purified recombinant ACO protein is visible as a single band around 35 KDa.

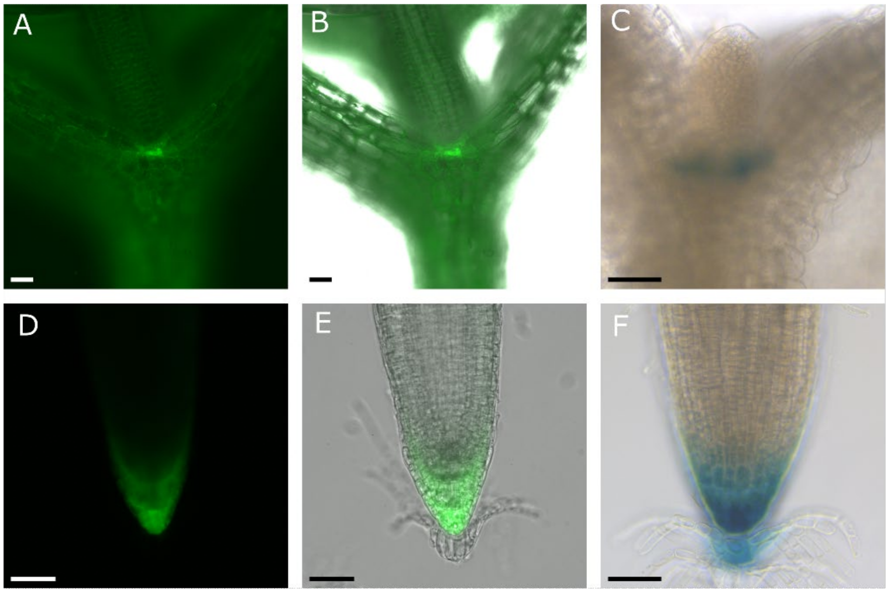


Supplemental Figure S5: *ACO1* expression is confined to the close proximity of apical meristems. The *pACO1::GFP-GUS* reporter shows a strong signal surrounding the shoot (A, B and C) and root apical meristem (D, E and F). GFP (A, B, D and E) and GUS (C and F) imaging showed a similar expression pattern. GFP was imaged (A and D) and overlaid with bright field images (B and E). Scale bars are 50 µm.

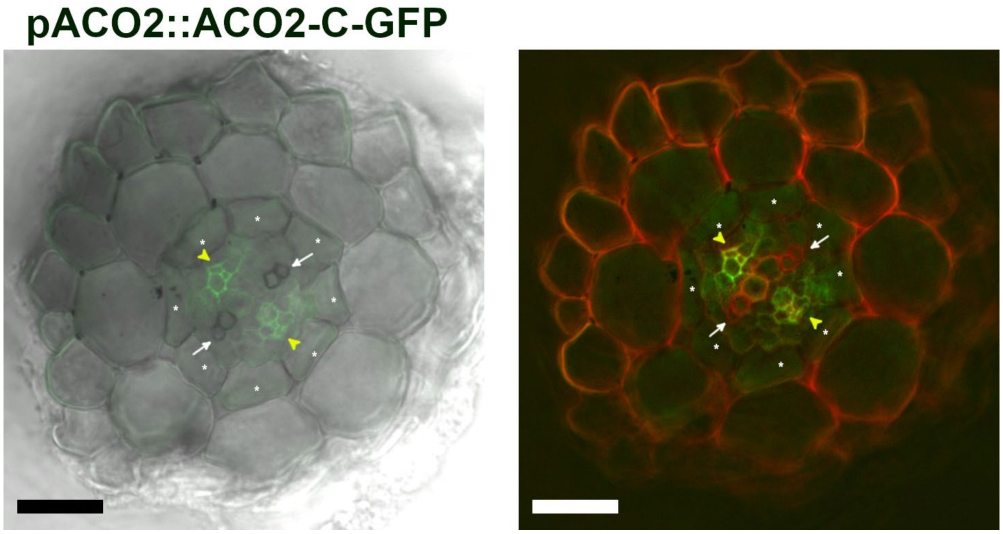


Supplemental Figure S6: ACO2 translational reporter shows phloem-specific localization. Primary roots of 7-day old plants were embedded in 3 % agarose and sectioned by hand. Sections were briefly stained with PI before imaging by fluorescence microscopy. (A) Overlay of brightfield and GFP channels and (B) overlay of PI and GFP channels. Asterisks indicate endodermal cells, white arrows indicate protoxylem, yellow arrowheads indicate phloem. Scale bars are 20 µm.

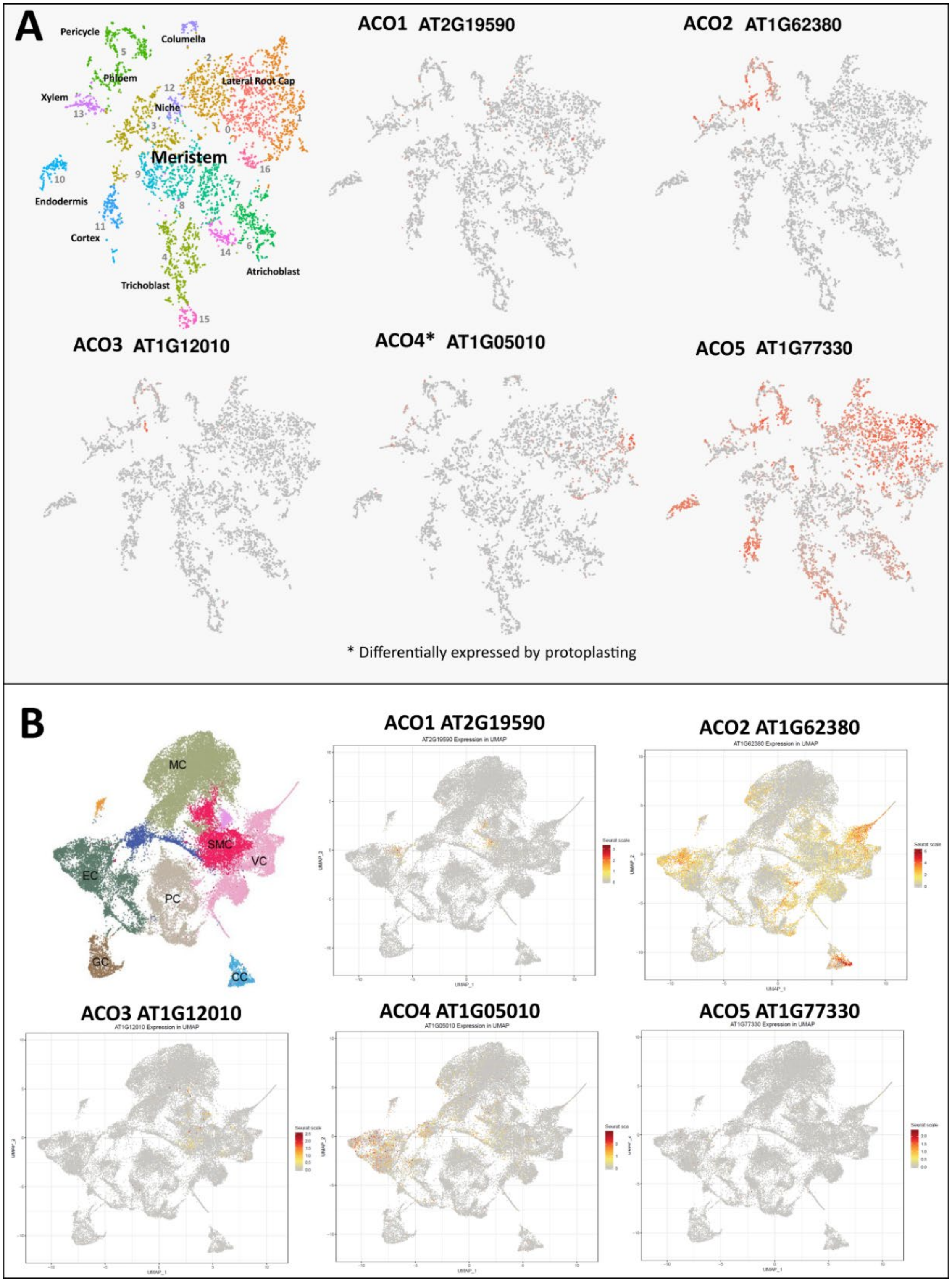


Supplemental Figure S7: Single-cell RNA-seq (scRNA-seq) data confirms *ACO* expression patterns observed in the *ACO* reporters. (A) Publicly available scRNA-seq data obtained from roots (Denyer at al., 2019; Ma et al., 2020) and (B) shoots (Zhang et al., 2021) with cell identity clusters visualized on the left. MC: mesophyll cell (olive); EC: endodermis cell (green); GC: guard cell (brown); PC: proliferating cell (gray); VC: vascular cell (pink); CC: companion cell (bleu); SMC: shoot meristem cell (purple).

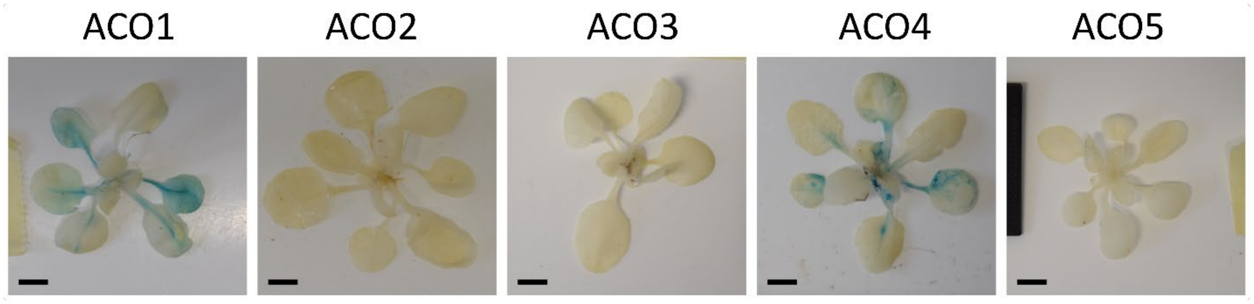


Supplemental Figure S8: ACOs show differential expression in rosette leaves. Rosette images of 3-week old p*ACO*$_{1-5}$:*GUS-GFP* transcriptional reporter lines. Scale bar equals 1 cm.

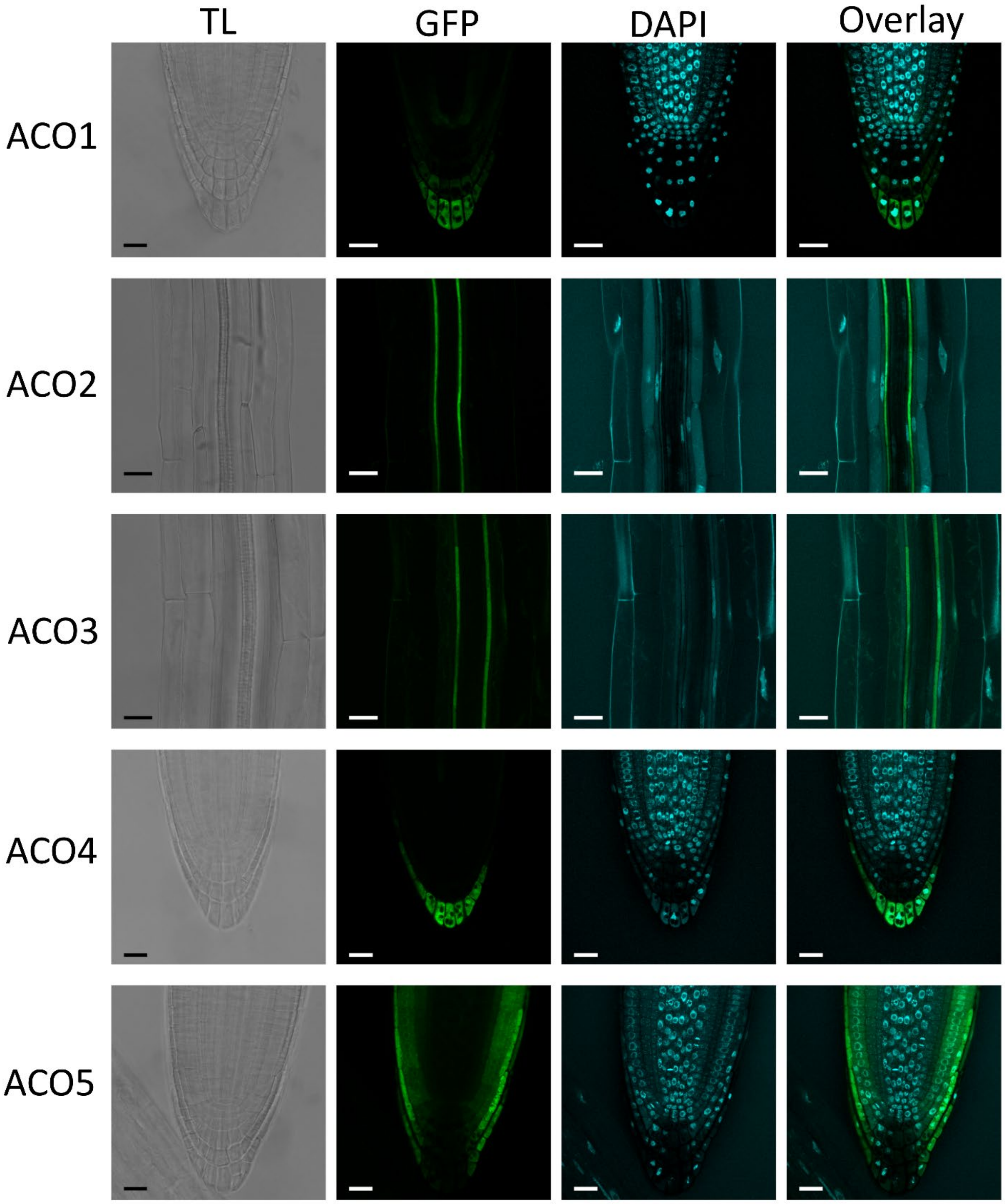


Supplemental Figure S9: ACOs show cytosolic and nuclear subcellular localization. Confocal microscopy of 5-day old translational p*ACO*$_{1\text{-}5}$:*GUS-GFP* reporter lines after fixation and clearing using ClearSee, and with DAPI staining. Rows show transmitted light (TL), GFP, DAPI, and an overlay of GFP and DAPI. Scale bars are 20 µm.

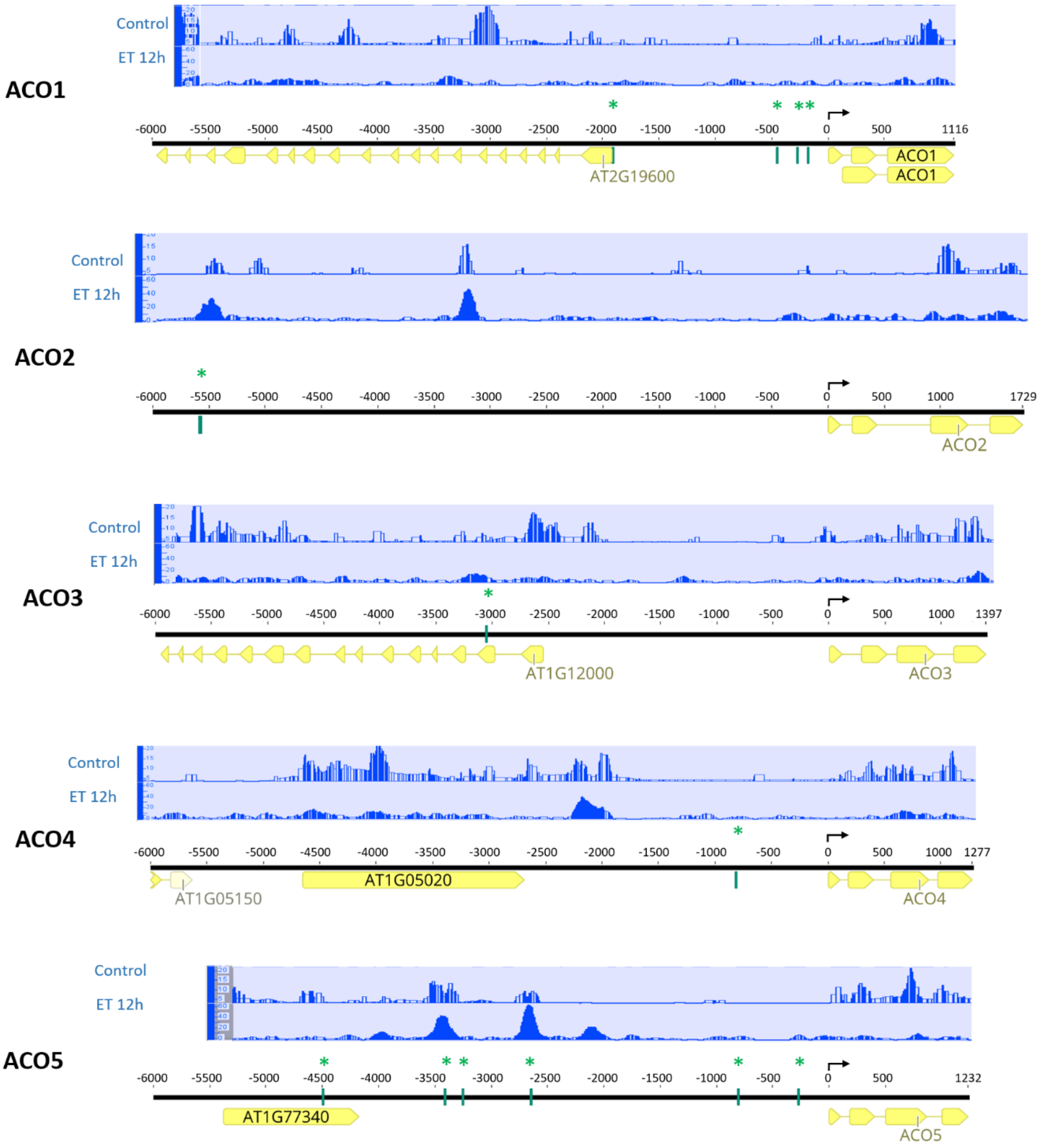


Supplemental Figure S10: The *ACO* promoters harbor EIN3 binding sites. Genomic regions of the *ACOs* showing their coding sequence (CDS) and a promotor region of 6 Kb upstream of the start codons (black arrow) and including other upstream CDS. EIN3 binding sites identified using FIMO (see methods) are indicated by green blocks and asterisks. Aligned ChIP-seq data (in the bleu zones above the gene models) showing EIN3 binding regions in control plants and plants treated with ethylene for 12 hours. Data was acquired from the Integrated Genome Browser (Freese et al., 2016).

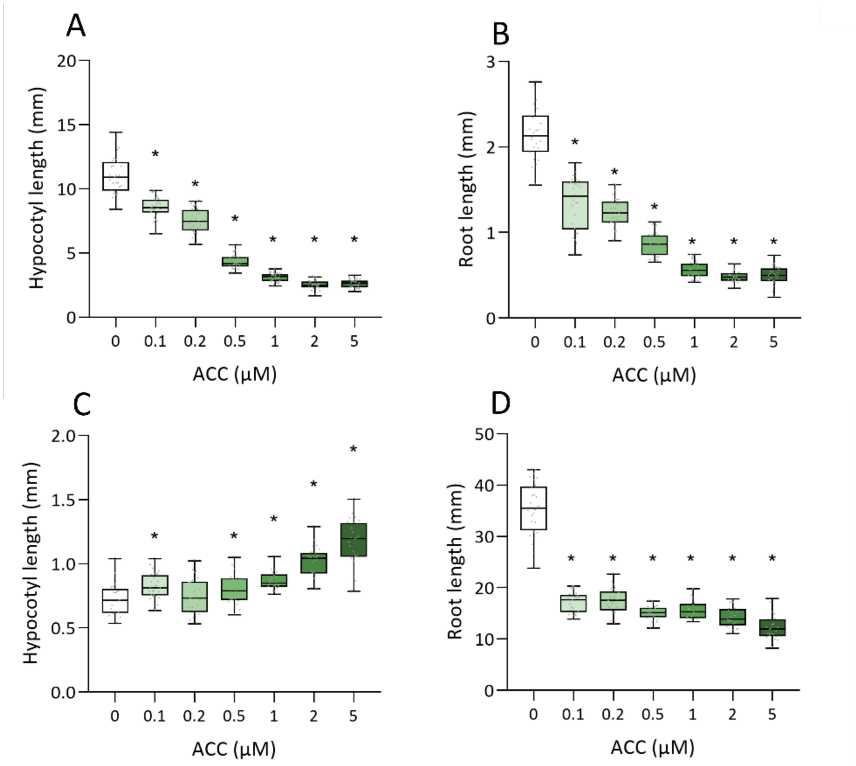


Supplemental Figure S11: Phenotyping of Col-0 wild type seedlings to an ACC gradient treatment. Hypocotyl (A and C) and root (B and D) lengths were measured (n ≥ 20) in 4-day old dark-grown (A and B) and 10-day old light-grown seedlings (C and D). Different shades of green depict different concentrations of ACC.One way ANOVA and Dunnett's post-hoc test were performed. Asterisks represent significance compared to Col-0 (p-value < 0.05).

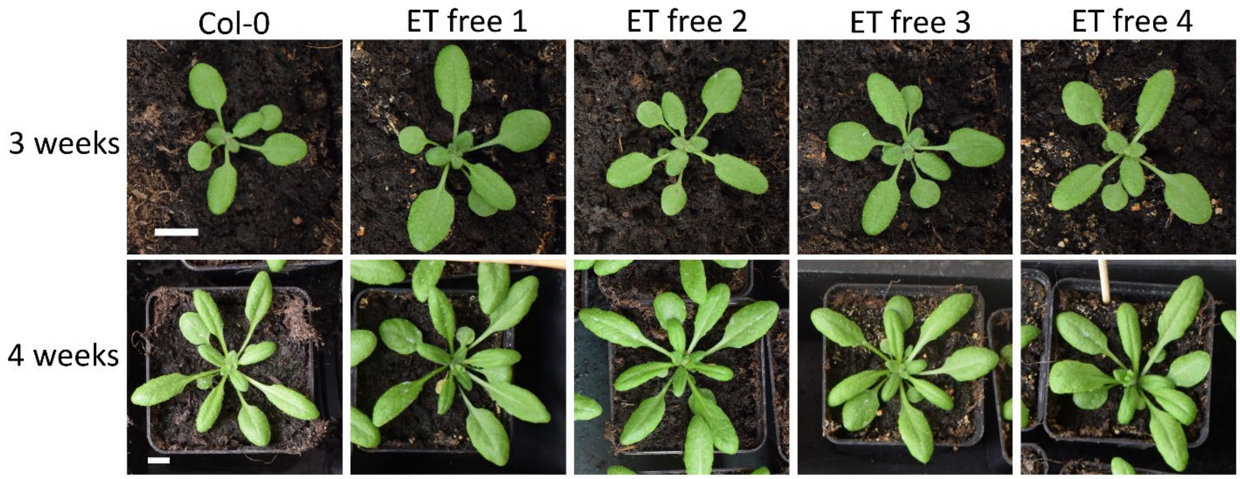


Supplemental Figure S12: Representative images of rosettes of 3- and 4-week-old plants of Col-0 and the different *aco* quintuple lines (ET free 1-4). Scale bar is 1 cm.

### Supplementary tables

Supplemental Table S1. Seed stocks used in this study.

| Name | Gene locus | SALK ID | NASC ID | Notes |
|---|---|---|---|---|
| ***aco1-1*** | AT2G19590 | SALK_127963 | N627963 | |
| ***aco2-1*** | AT1G62380 | SALK_027311 | N527311 | |
| ***aco2-2*** | AT1G62380 | GABI_696A02 | NA | *aco2-2* introduced into higher order mutants by crossing with *aco2,3,4* mutant |
| ***aco3-1*** | AT1G12010 | SALK_082132 | N582132 | |
| ***aco4-1*** | AT1G05010 | SALK_014965 | N514965 | |
| ***aco5-1*** | AT1G77330 | SAIL_759_D06 | N876603 | Only used to generate translational reporter, all phenotyping experiments were conducted in *aco5-2* |
| ***aco5-2*** | AT1G77330 | SALK_042400 | N542400 | |

Supplemental Table S2. Primers used in this study.

| Experiment | Primer name | Primer sequence 5'-3' | Notes |
|---|---|---|---|
| **Genotyping primers** | aco1-1_F | CTTTCCCAATCTGCATCTGAG | |
| | aco1-1_R | AACACAACACCACGCATATCC | |
| | aco2-1_F | CAGATCGGTTTAGCTGTACCG | |
| | aco2-1_R | TCGAGAAGATGACAAAGGACC | |
| | aco2-2_F | CATGGTGGATAATTGCTCACC | |
| | aco2-2_R | TTGCGCCTATAAATAGCAACC | |
| | aco3-1_F | ATCCCATCTCAAAAGCAGGAG | |
| | aco3-1_R | CTTGAAACAGCAAAATGAGGC | |
| | aco4-1_F | GTCCATATGCATTTGGACTGG | |
| | aco4-1_R | GGAGCTACTGGATCTGCTGTG | |
| | aco5-2_F | TCAGGGTCCATAAGAATGACG | |
| | aco5-2_R | TATGATGGCCTTCAGGTCTTG | |
| | Salk_LBb1.3 | ATTTTGCCGATTTCGGAAC | |
| **CRISPR primers** | ACO5_sgRNA_F | TTTTGGTCTCAATTGgcgctagagcttgcgaagagGTTTTAG AGCTAGAAATAGC | Lower case indicates sgRNA target site |
| | ACO5_sgRNA_R | TTTTGGTCTCAAAACtcatcttgctcgctagcttcCAATCACT ACTTCGACTC | Lower case indicates sgRNA target site |
| **Transcriptional reporters** | pAtACO1_F | ggggacaagtttgtacaaaaaagcaggctCGCCTAATGAGCAC AACACA | Lower case indicates GW sites |
| | pAtACO1_R | ggggaccactttgtacaagaaagctgggtAGCCTTGACCATCTC TGACT | Lower case indicates GW sites |
| | pAtACO2_F | ggggacaagtttgtacaaaaaagcaggctACACCGTGCTTGCA CTCTTT | Lower case indicates GW sites |
| | pAtACO2_R | ggggaccactttgtacaagaaagctgggtCCTTTGGACTTGAGC ATGTCA | Lower case indicates GW sites |
| | pAtACO3_F | ggggacaagtttgtacaaaaaagcaggctCTCTCTCTCTCTCTC TTAACTAGC | Lower case indicates GW sites |
| | pAtACO3_R | ggggaccactttgtacaagaaagctgggtTTCAACTTCGGTCTC GAGGG | Lower case indicates GW sites |
| | pAtACO4_F | ggggacaagtttgtacaaaaaagcaggctCTCTCTCTCTTTTTTT TTAAATGGGTTTC | Lower case indicates GW sites |

| | | | |
|---|---|---|---|
| | pAtACO4_R | ggggaccactttgtacaagaaagctgggtCTCCACTTTGTCCAAAAGCTC | Lower case indicates GW sites |
| | pAtACO5_F | ggggacaagtttgtacaaaaaagcaggctACTTAAAAGTGATCATTTCCTTATGG | Lower case indicates GW sites |
| | pAtACO5_R | ggggaccactttgtacaagaaagctgggtACGTTTTCTAGCTTCTCGCCA | Lower case indicates GW sites |
| **Translational reporter** | AtACO1_prom_pGGAf | AACAggtctcAACCTTTGCAACAACCAACCTG | For cloning to pGGA000, lower case indicates BsaI site |
| | AtACO1_prom_pGGAr | AACAggtctcATGTTCTCTTTTTTATTTACTTTTTCTCAC | For cloning to pGGA000, lower case indicates BsaI site |
| | AtACO1_CDS-stop_pGGCf | AACAggtctcAGGCTCGATGGTTTTGATCAAAGAGAGAGA | For cloning to pGGC000, lower case indicates BsaI site |
| | AtACO1_CDS-stop_pGGCr | AACAggtctcACTGAGGCTGAATCCGCATTTCCCATT | For cloning to pGGC000, lower case indicates BsaI site |
| | AtACO2_prom_pGGAf | AACAggtctcAACCTCACCGTGCTTGCACTCTTT | For cloning to pGGA000, lower case indicates BsaI site |
| | AtACO2_prom_pGGAr | AACAggtctcATGTTCTTTCTTTCTCTCTCTCTTCTTTGA | For cloning to pGGA000, lower case indicates BsaI site |
| | AtACO2_CDS-stop_pGGCf | AACAggtctcAGGCTCGATGGAGAAGAACATGAAGTTTCC | For cloning to pGGC000, lower case indicates BsaI site |
| | AtACO2_CDS-stop_pGGCr | AACAggtctcACTGAGAAAGTCTCTACGGCTGCTG | For cloning to pGGC000, lower case indicates BsaI site |
| | AtACO3_prom_pGGAf | AACAggtctcAACCTCGCATTGGGGTTGGTGT | For cloning to pGGA000, lower case indicates BsaI site |
| | AtACO3_prom_pGGAr | AACAggtctcATGTTCTCTCTCTCTCTCTCTTAACTAGC | For cloning to pGGA000, lower case indicates BsaI site |
| | AtACO3_CDS-stop_pGGCf | AACAggtctcAGGCTCGATGGAGATGAACATTAAGTTTCC | For cloning to pGGC000, lower case indicates BsaI site |
| | AtACO3_CDS-stop_pGGCr | AACAggtctcACTGAGAATGTCTCAACCACAGCC | For cloning to pGGC000, lower case indicates BsaI site |
| | AtACO4_prom_pGGAf | AACAggtctcAACCTCCCATTGGCCATTGACTA | For cloning to pGGA000, lower case indicates BsaI site |
| | AtACO4_prom_pGGAr | AACAggtctcATGTTCTCTCTCTCTTTTTTTTTAAATGGGTTTC | For cloning to pGGA000, lower case indicates BsaI site |
| | AtACO4_CDS-stop_pGGCf | AACAggtctcAGGCTCGATGGAGAGTTTCCCGATCATC | For cloning to pGGC000, lower case indicates BsaI site |
| | AtACO4_CDS-stop_pGGCr | AACAggtctcACTGACGCAGTGGCCAATGGTC | For cloning to pGGC000, lower case indicates BsaI site |
| | AtACO5_prom_pGGAf | AACAggtctcCAACCTAGCAGGCTGTGATCATTTCC | For cloning to pGGA000, lower case indicates BsaI site |
| | AtACO5_prom_pGGAr | AACAggtctcAACCTACTTAAAAGTGATCATTTCCTTATGG | For cloning to pGGA000, lower case indicates BsaI site |
| | AtACO5_CDS-stop_pGGCf | AACAggtctcAGGCTCGATGGCGATTCCTGTTATCGATTTC | For cloning to pGGC000, lower case indicates BsaI site |
| | AtACO5_CDS-stop_pGGCr | AACAggtctcACTGAGAGAGACTTTACAGCTAGAAAACG | For cloning to pGGC000, lower case indicates BsaI site |
| **qRT-PCR** | AtACO1_FW_Q2 | ATACCCAGAATGCCCACGTC | |
| | AtACO1_RV_Q2 | GGATGGCGGTATAGGAACCC | |
| | AtACO2_FW_Q2 | AGATGTCGATTGGGAAAGCA | |
| | AtACO2_RV_Q2 | TCCAACAAATCCTCAGCAAG | |
| | At_ACO3Q_FW | GAAATGCTTCGTTCCAAAGG | |
| | At_ACO3Q_RV | GCCTCTTCCCAAAATCCTTC | |
| | At_ACO4Q_FW | CCGGTTAAGCATTCAATCGT | |
| | At_ACO4Q_RV | AGAGTCGCTTCCCGGATTAT | |
| | At_QRT_ACO5_RV | GCTAGAAAACGAGGCTCTTTAGG | |
| | At_QRT_ACO5_FW | GGGAGGAAGGAAACAGAAGG | |
